# *Cis*-regulatory chromatin loops arise before TADs and gene activation, and are independent of cell fate during development

**DOI:** 10.1101/2020.07.07.191015

**Authors:** Sergio Martin Espinola, Markus Götz, Jean-Bernard Fiche, Maelle Bellec, Christophe Houbron, Andrés M. Cardozo Gizzi, Mounia Lagha, Marcelo Nollmann

**Author notes:** Co-first authors (Sergio Martin Espinola, Markus Götz).

## Abstract

During development, naïve cells gradually acquire distinct cell fates, through sophisticated mechanisms of precise spatio-temporal gene regulation. Acquisition of cell fate is thought to rely on the specific interaction of remote *cis*-regulatory modules (e.g. enhancers, silencers) (CRM) and target promoters. However, the precise interplay between chromatin structure and gene expression is still unclear, particularly in single cells within multicellular developing organisms. Here we employ Hi-M, a single-cell spatial genomics approach, to systematically detect CRM-promoter looping interactions within topological associating domains (TADs) during *Drosophila* development. By comparing *cis*-regulatory loops in alternate cell types, we show that physical proximity does not necessarily instruct transcriptional states. Moreover, multi-way analyses revealed the existence of local interactions between multiple remote CRMs to form hubs. We found that loops and CRM hubs are established early during development, prior to the emergence of TADs. Moreover, CRM hubs are formed via the action of the pioneer transcription factor Zelda and precede transcriptional activation. Our approach offers a new perspective on the role of CRM-promoter interactions in defining transcriptional activation and repression states, as well as distinct cell types.

## Introduction

Chromosomes are organized at different levels —nucleosomes, chromatin loops, topologically associating domains (TADs) and chromosome territories— and each of these layers contributes to the regulation of transcription ^1,2^. For instance, post-translational histone modifications are essential for bookmarking transcriptional states and epigenetic memory at the nucleosomal scale, while local loops between enhancers (E) and promoters (P) are critical for the precise regulation of transcriptional activation ^3–7^. In addition, organization of chromosomes into TADs plays a role in transcriptional regulation ^8^, primarily by facilitating communication between enhancers and promoters through E-P loops within a TAD and restricting contacts from enhancers of neighboring TADs ^5,9–13^. However, the interplay between formation of E-P loops, emergence of TADs, and transcriptional output is still poorly understood ^14^.

Tissue-specific enhancers have been shown to be in close proximity with their cognate promoters indicating that E-P contacts are needed for precise gene regulation ^15–18^. Indeed, introduction of ectopically-enforced enhancer-promoter contacts lead to transcription activation of a reporter gene during *Drosophila* development ^19^. In some cases, enhancers can increase transcriptional output by modulating transcriptional bursting frequency ^6,7,20–22^. However, in other cases E-P contacts seem to be dissociated from gene activation. For instance, in mice gene activation was observed without E-P proximity ^23,24^, suggesting that an enhancer may not necessarily need to be in continuous physical contact with a promoter to influence transcription. The mechanisms by which E-P contacts may regulate transcription are currently under intense debate ^14,25,26^.

Early evidence showed that promoters can contact several distant enhancers ^15–17^, raising the possibility that more than one enhancer may contact a promoter at any given time. More recently, use of multi-way 3C and 4C methods showed that, indeed, enhancers can cluster together to form enhancer hubs that can recruit one or several promoters ^27–30^. This is supported by evidence of nuclear microenvironments containing multiple enhancers and clusters of transcription factors ^31–37^. This model is consistent with multi-way interactions between distal enhancers to regulate promoter activity of single or multiple genes by sharing resources. Whether and how formation of multi-way interactions may be related to the emergence of TADs during development ^38,39^ is still an open question.

To shed light into these questions, we investigated the interplay between transcriptional state and physical proximity between promoters and large sets of *cis*-regulatory modules (CRMs, e.g. enhancers, silencers and insulators) during the awakening of the zygotic genome in early *Drosophila* embryos. During the first hours of development, *Drosophila* embryos offer an ideal biological context to decipher how CRMs are employed to establish precise spatio-temporal patterns of gene expression. Decades of genetic and genomic studies have characterized CRMs at a large scale and their usage to interpret morphogen gradients ^40–42^. In particular, the pioneering activity of factors such as Zelda (Zld) establish early accessibility of CRMs (reviewed in ^43^) and is involved in the emergence of TAD organization ^38,44^.

Here, we used Hi-M, an imaging-based technology enabling the detection of chromatin organization and transcriptional status in intact embryos ^45,46^. This technology allowed us to visualize where and when interactions between CRMs occur and question their impact on transcriptional states. We first used Hi-M to detect intra-TAD chromatin loops in *Drosophila* embryos. We show that the majority of these loops involve CRMs. In fact, we identified not only E-P loops but multiple CRM contacts (E-P, P-P and E-E) co-interacting locally in single cells and referred to as CRM hubs. Unexpectedly, these contacts were not found to be specific to transcriptionally active cells. Hence, tissues with different cell fates, exhibit similar CRMs contacts and E-P loops. Moreover, networks of CRM loops are established at early stages, prior to the emergence of TADs and before transcriptional activation. Finally, we provide evidence that the pioneer factor Zld is required for the establishment of subsets of CRM hubs.

## Results

### High-resolution Hi-M reveals preferential interactions between *cis*-regulatory modules

Functional characterization of specific chromatin loops between *cis*-regulatory modules within TADs (Fig. 1a) requires the development of technologies adapted for the simultaneous detection of such looping interactions and of transcriptional output. Recently, we and others established a new family of imaging-based methods able to retrieve chromatin architecture and transcriptional status simultaneously in single cells (Hi-M and optical reconstruction chromatin architecture, or ORCA) ^45–47^. Hi-M relies on the labeling and imaging of the expression pattern of genes by direct detection of transcripts via RNA-FISH, followed by the sequential imaging of tens of distinct DNA loci by oligopaint-FISH ^48^ in intact *Drosophila* embryos ^45,46^. First, we tested whether conventional Hi-M was able to detect intra-TAD chromatin loops. For this, we took advantage of the numerous whole genome profiling datasets available for the early *Drosophila* embryo to select two regions harbouring early developmental genes expressed at different timings and in different regions of the embryo (*dorsocross* (*doc*)- and *snail* (*sna*)*-*TADs).

**Figure 1.**
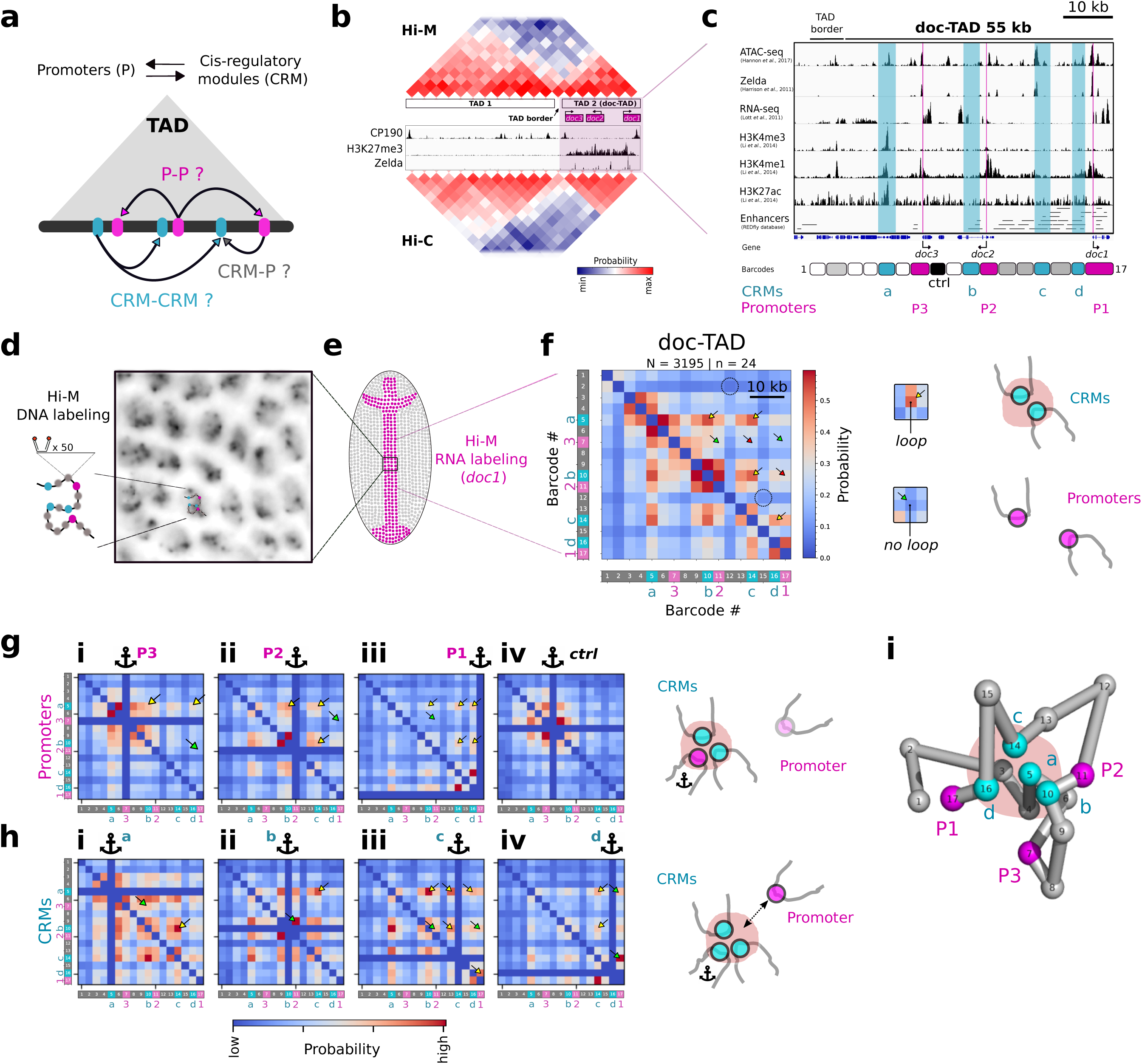
Hi-M reveals widespread *cis*-regulatory chromatin loops and hubs within TADs. a. A main aim of the study is to investigate the networks of contacts between *cis*-regulatory modules (CRMs, cyan) and promoters (P, magenta) within TADs (gray triangle). Hypothetical chromatin loops are shown by arrows. b. The *doc* locus (Chr3L:8.88..9.03Mb) in *Drosophila melanogaster*. Low-resolution Hi-M and Hi-C ^44^ contact probability maps are shown on top and bottom, respectively. Blue and red indicate low and high contact probabilities, respectively. Two TADs can be clearly distinguished: TAD 2 comprises the three *doc* genes (*doc*-TAD, shaded region) and is flanked by insulator binding sites (CP190), displays polycomb marks (H3K27me3), as well as several Zelda peaks. c. Epigenetic profile of the *doc*-TAD. Tracks for chromatin accessibility (ATAC-seq), pioneer factor binding (Zelda), transcriptional activity (RNA-seq), chromatin marks for active promoters (H3K4me3), and active enhancers (H3K4me1, H3K27ac), and RedFly enhancers are shown. Lower panel shows the barcodes used for high-resolution Hi-M: barcodes with enhancer marks are annotated as CRMs (a-d, cyan), with *doc* promoters are annotated as P1-P3 (magenta), a control barcode is annotated as ‘ctrl’ (black), and barcodes with enhancer marks annotated in gray. See Supplementary Table 1 for assignment of CRM_b-d_. d. Schematic diagram of the labeling strategy. Left: Each barcode is composed of 50 assemblies of a primary oligonucleotide containing a homology region (black) and two fluorescent readout probes (orange). Right: Nuclei are imaged by DAPI staining (dark gray). The physical location of each barcode inside the nucleus is reconstructed with nanometric precision from multiplexed, sequential imaging. e. Schematic representation of a *Drosophila* embryo (dorsally oriented). Segmentation of actively transcribing cells (magenta dots) is based on nascent RNA FISH labeling in whole mount embryos. f. The high-resolution Hi-M contact probability map of the *doc*-TAD derived from cells displaying *doc1* expression in nc14 embryos. The contact probability is color-coded according to the colorbar on the right. Barcodes are indicated on the bottom and left axis with cyan for CRMs and magenta for promoters. Yellow arrows indicate strong looping interactions between CRMs. Green arrows represent the bins where promoter-promoter interactions are expected. Red arrows show examples of bins with a CRM and a promoter. Black circles represent regions with predicted enhancers (RedFly) or with active chromatin (ATAC-seq peak) that do not display looping interactions. Insets show a region of the matrix with a preferential looping interaction (*loop*) as well as a region without any preferential loop detected (*no loop*). The scheme to the right illustrates the chromatin organization of CRMs and promoters. Neighboring barcodes often displayed high interaction frequencies due to the polymer nature of the chromatin fiber, therefore we do not consider them here as specific chromatin loops. Number of nuclei with *doc1* expression: N=3195. Number of embryos with *doc1* expression: n=24. Total number of examined nuclei: 37129, total number of embryos: 29. g. Multi-way interactions between promoter regions. The 3-way contact probability is color-coded according to the colorbar. Anchoring barcodes are highlighted by a pictogram and placed at the three promoter regions (panels i-iii) or the control barcode (panel iv). Barcodes are indicated on the left and bottom axis as in panel f. Prominent peaks (yellow arrows) comprise one promoter and two CRMs but not multiple promoters (green arrows). The scheme to the right illustrates the spatial arrangement of CRMs and promoter regions when the anchor is placed at a promoter. h. Multi-way interactions between CRMs. Color-code as in panel g. Anchors are placed at the four CRMs. Prominent peaks are found for a combination of three CRMs (yellow arrows) and with lower probability for a combination of two CRMs and a promoter region. The scheme to the right illustrates the spatial arrangement of CRMs and promoter regions when the anchor is placed at a CRM. i. 3D topological reconstruction of the *doc*-TAD. CRMs and promoter regions are indicated as cyan and magenta spheres, respectively. CRMs tend to cluster in the center of the TAD while promoter regions are found more likely at the periphery.

The *doc*-TAD contains a family of three genes, the *dorsocross* genes *doc1, doc2*, and *doc3* encoding T-box transcription factors. These genes display similar expression patterns, particularly during early stages of embryogenesis, in the rapidly-developing pre-gastrulation blastoderm embryo (nuclear cycle, nc 11 to 14), which will be the focus of this study (Figs. S1a-b). Genetic studies revealed that these genes are functionally redundant and are essential for the development of the amnioserosa and cardiogenesis ^49^. In early embryos, the *doc*-TAD is flanked by insulator binding sites (e.g. CP190), and displays extensive H3K27me3 marks as well as several prominent Zelda peaks (Fig. 1b) ^50–53^. At nc14, the Hi-M contact probability map of this genomic region displays two clear TADs, similar to those detected by Hi-C (TAD1, and *doc*-TAD, Figs. 1b, S1a) ^44^. Inspection of ATAC-seq ^54^, H3K4me3, H3K4me1, and H3K27ac profiles^55^ as well as of enhancer databases ^56^ revealed that the *doc*-TAD contains several putative CRMs, including four potential enhancers (CRM_a_, CRM_b_, CRM_c_ and CRM_d_) for the three *doc* promoters (Fig. 1c, Supplementary Table 1) ^50,51,56,57^. However, conventional Hi-M/Hi-C did not exhibit specific looping interactions within the *doc*-TAD, most likely due to insufficient genomic resolution and coverage (Fig. 1b).

To overcome these limitations and probe communications between CRMs and promoters within TADs in an unbiased manner, we improved the genomic resolution and coverage of Hi-M by 3-fold (from ∼8-10kb to ∼3kb) and painted the entire *doc*-TAD with contiguous barcodes, particularly targeting promoters and predicted CRMs (Figs. 1c, S1a). We first focused on enhancers already validated by transgenic assays (CRM_b-d_) (Supplementary Table 1). However, we note that other genomic regions in this TAD (e.g. CRM_a_) contain characteristic enhancer marks but are not present in enhancer databases (Figs. 1c, and Supplementary Table 1). The three *doc* genes within the *doc*-TAD exhibit a shared spatio-temporal profile of expression in late nc14 (Fig. S1b). Thus, we hypothesized that multiple putative CRMs are likely to contact *doc* promoters to regulate their common expression pattern.

To test this hypothesis, we obtained contact maps of transcriptionally active cells by combining high-resolution Hi-M to nascent mRNA-FISH labeling (Fig. 1d-f). Remarkably, the improvement in genomic coverage in Hi-M now enabled the detection of specific looping interactions between genetic elements within the *doc*-TAD in intact embryos (Figs. 1f, S1h). The strongest loops represented in all cases interactions between CRMs (Fig. 1f, yellow arrows). To quantify the strength of looping interactions, we generated virtual interaction profiles from Hi-M data (hereafter 4M plots) where we mapped the contact probabilities of a single predefined anchor with the rest of the barcodes. For instance, CRM_c_ predominantly interacts with CRM_a_ and CRM_b_ with similar probabilities (Fig. S1c-vii). In contrast, we did not observe specific loops between CRMs and barcodes not containing CRMs (e.g. ctrl barcode, Figs. 1c and S1c-iv). Interactions between CRMs and promoter regions (e.g. P1, P2 and P3) were present but displayed lower frequencies than interactions between CRMs (red arrows, Fig. 1f). Finally, contacts between promoters were in all cases rather weak (green arrows, Figs. 1f, S1c-i,ii,iii).

Next, we investigated whether all putative CRMs displayed chromatin loops. Interestingly, a CRM predicted from epigenetic profiling but not present in enhancer databases (e.g. CRM_a_, see complete list of reported enhancers in Supplementary Table 1) displayed extensive interactions with reported enhancers (e.g. CRM_b_, CRM_c_, CRM_d_) as well as with the promoters of *doc* genes (Fig. S1c-v). In contrast, a subset of barcodes harbouring previously described enhancers, or displaying enhancer marks (e.g. ATAC-seq, H3K4me1, see barcodes 2, 12, 13, 15 in Fig. 1C) failed to exhibit specific looping interactions with other CRMs (e.g. black circles in Fig. 1f), perhaps because they are not active in dorsal cells at nc14. Thus, high-resolution Hi-M reveals unforeseen interactions between heretofore unreported CRMs and other regulatory regions within the *doc*-TAD, and permits the quantification of the frequencies with which putative enhancers actively contact cognate target promoters in a specific tissue and developmental time.

Encouraged by these results, we applied a similar procedure to the *sna*-TAD, containing a family of paralogous genes encoding the zinc finger transcription factor *snail (sna)*, in addition to *worniu (wor)* and *escargot (esg)* genes, as well as multiple CRMs (Fig. S1d). Remarkably, loops between CRMs were also highly common in this locus, particularly between known enhancers of *sna* and of *esg* (Figs. S1d-e, Supplementary Table 1). Collectively, these data suggest that promoters interact with a panoply of enhancers that can be shared between the different genes within a TAD.

### Shared enhancers, promoter competition and CRM hubs

The existence of multiple pairwise interactions between CRMs within the *doc*-TAD and the naturally-occurring overlapping expression patterns of *doc* genes (Fig. S1b) suggests that multiple CRMs may compete or cooperate for gene activation in single cells. To discriminate between these two hypotheses, we tested whether multi-way interactions are formed by excluding an anchor of interest and plotting the frequencies with which two barcodes interact together with this given anchor ^28^. First, we selected promoters as anchors. We observed that promoters do not tend to contact other promoters (green arrows, Fig. 1g-i,ii,iii), consistent with our previous observations from Hi-M contact maps (Figs. 1f and S1c). Instead, the three *doc* promoters preferentially looped to multiple CRMs in single cells (yellow arrows, Fig. 1g-i,ii,iii). A control locus with no promoter marks failed to display specific looping interactions (Fig. 1g-iv). Overall, these data support a promoter competition model, as recently proposed in *Escherichia coli* ^58^.

Genomic methods revealed the spatial clustering of multiple enhancers in cultured mammalian cells ^27,28^. To test whether spatial clustering of multiple enhancers could be directly visualized in intact embryos, we mapped 3-way interactions using CRMs as anchors. Interestingly, we observed that CRM_a-d_ displayed high frequencies of multi-way interactions (Fig. 1h, see examples labeled by yellow arrows). By analogy to what has been described in cultured cells using genome-wide methods ^27,28^, we termed these CRM interaction networks “CRM hubs”. CRM hubs can contain promoters (green arrows, Fig. 1h), but most often contained known or putative enhancers. Analysis of the *sna*-TAD reveals a similar scenario, where mostly barcodes containing CRMs are involved in most 3-way interactions (see for example interactions between *esg* and *sna* enhancers in Figs. S1g-iv, v and vi).

To confirm the existence of CRM hubs with an independent analysis approach we turned to the use of ShRec3D, a method that enables the conversion of contact matrices into topological representations ^59^. We applied ShRec3D to the Hi-M map of *doc*-TAD, and obtained a topological reconstruction where CRMs can be clearly seen to cluster at the center of the TAD whereas promoter elements tend to be positioned at the periphery (Fig. 1i). Similarly, we observed that CRMs within *sna*-TAD also tended to cluster together at the center of the TAD (Fig. S1f). Collectively, 3-way and topological analyses suggest that multiple enhancers physically interact in a local nuclear space to form CRM hubs.

Critically, CRM hubs can but do not tend to contain multiple promoters. For example, the putative shared enhancer CRM_c_, located at 10 kb and 11 kb from *doc1* and d*oc2* TSS respectively, is contacted in most cells by other CRMs (yellow arrows, Fig. 1h-iii) and is contacted by multiple simultaneous promoters less often (green arrows, Fig. 1h-iii). These observations are consistent with our previous analyses showing that formation of promoter-promoter loops and promoter clusters is uncommon (Figs. 1f, 1g and S1c). Taken together, our results are inconsistent with: (a) promoters forming mutually-exclusive interactions with enhancers; and (2) multiple promoters coming together in space to share a common enhancer. Instead, our data suggest a model in which promoters contact CRM hubs containing multiple enhancers, as observed recently in bacteria and in mouse cells ^29,58^.

### Networks of long-range CRM contacts are indistinguishable between cells of different transcriptional status or cell fates

Next, we examined whether loops between CRMs depended on transcriptional status (repression/activation). For this, we investigated chromatin organization by Hi-M in three populations of cells emanating from the three main presumptive tissues established along the dorso-ventral axis during nc14: mesoderm (M), neuroectoderm (NE) and dorsal ectoderm (DE) ^60^. To distinguish between these three cell fates, we employed double RNA-FISH labeling prior to Hi-M (with *sna* and *doc* probes directly labeling mesodermal and dorsal ectodermal cells, respectively, Figs. 2a). Cells were classified as: (a) dorsal ectoderm cells when an active *doc1* transcription hotspot could be visualized (Fig. S1b); (b) mesoderm cells when located within the *sna* expression pattern (Fig. 2a); (c) neuroectoderm cells when located between the pattern of *sna* and the edge of the *doc1* pattern (Fig. 2a).

**Figure 2.**
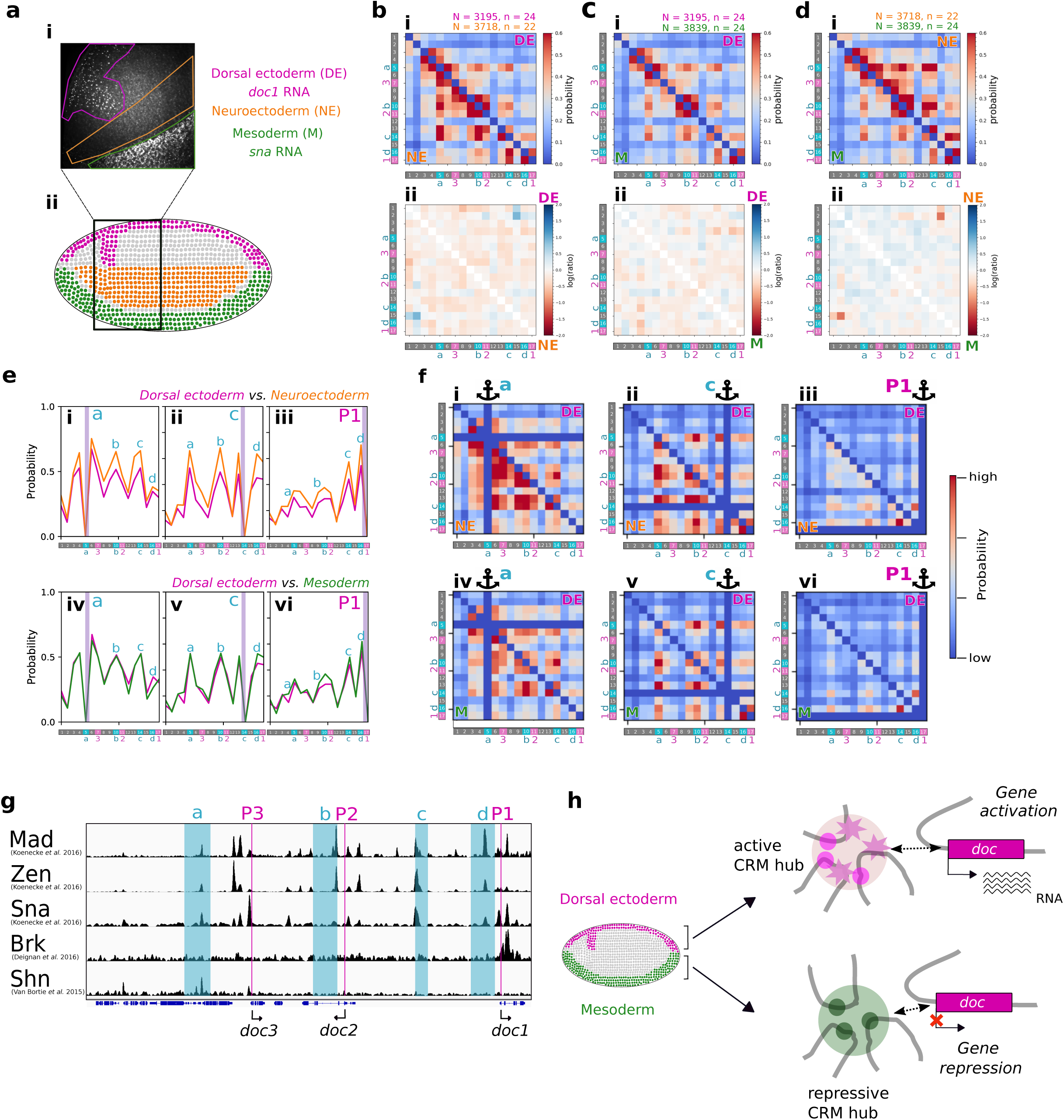
CRM-CRM and CRM-P loop frequencies are similar between cell types. a. Scheme indicating the three presumptive tissues and their segmentation (panel ii) based on double RNA-FISH labeling (panel i). Dorsal ectoderm (DE) is highlighted in magenta, neuroectoderm (NE) in orange, and mesoderm (M) in green. b. Contact probability maps for DE (upper-right half) and NE (lower-left half) (panel i). The map of the natural log of the ratio between the contact probabilities of DE and NE (panel ii). Blue indicates a larger contact probability in DE, red in NE. c. Similar to panel b, but for DE and M. d. Similar to panel b, but for NE and M. e. 4M profiles derived from Hi-M maps for a selected number of anchors. Anchors (indicated by vertical purple lines) were placed at CRM_a_ (panels i, iv), CRM_c_ (panels ii, v), and P1 (panels iii,vi). Upper row shows comparison between DE and NE, whereas the lower row presents the comparison between DE and M. Peaks are labeled with the corresponding CRM (see barcodes in the x-axis). f. Comparison of 3-way contacts for the same tissues and anchors as in panel e. The 3-way contact probability is color-coded according to the colorbar. Anchors are indicated by a pictogram. Barcodes are shown at the bottom and left of each map. g. ChIP profiles of key transcriptional regulators in the *doc*-TAD. Activators (Mad, Zen) and repressors (Brk, Sna, Shn) both bind CRMs. h. Illustration of the double role of CRMs in the *doc*-TAD: Activators (magenta star and circle) form active CRM hubs in the dorsal ectoderm and can reinforce transcription when contacting *doc* genes (RNA represented as wavy lines). In contrast, repressors (green circles) bind CRMs in the mesoderm and neuroectoderm and can silence the expression of *doc* genes.

Unexpectedly, Hi-M interaction matrices for DE, NE and M displayed only minor differences (Figs 2b-2d), indicating that the same network of CRM loops is present in cells that are actively transcribing and in cells that are programmed to repress expression of *doc* genes at this nuclear stage. To analyze these looping interactions quantitatively, we extracted 4M profiles from Hi-M matrices. The 4M profiles were almost identical in cells with different cell fates and activation status, independently of whether promoters or CRMs were used as anchors (Figs. 2e, S2a-b). For instance, the *doc1* promoter (P1) showed identical interactions with the four CRMs (CRM_a-d_) in the dorsal-ectoderm, the neuroectoderm and the mesoderm (Fig. 2e-iii,vi). Likewise, CRM_a_ and CRM_c_ displayed patterns of interactions with other CRMs that were indistinguishable between tissues (Fig. 2e-i,ii,iv,v).

Finally, to detect whether CRM hubs existed in tissues where *doc* genes are repressed, we performed single-cell 3-way analyses. Indeed, comparison of 3-way interaction matrices of NE and M with those of DE revealed the persistence of CRM hubs in cells where transcription is repressed (Fig. 2f), suggesting that CRM hubs also exist in these cell types.

To test whether this surprising similarity between CRM loops in alternative cell fates could be observed in other genomic regions, we explored chromatin organization in the *sna* locus. We obtained similar networks of 2-way and 3-way interactions in mesoderm cells versus cells in other tissues (neuro and dorsal-ectoderms, Fig. S2c-f). The major differences in Hi-M contact maps between tissues in the *sna*-TAD arose from a depletion of interactions in the mesoderm with respect to those in the dorsal- and neuro-ectoderms (see red stripes, Fig. S2c). However, local interactions networks around promoters (e.g. contacts between *sna* and its shadow enhancer, insets of Fig. S2c) as well as long-range interactions between enhancers (e.g. between *sna* and *esg* enhancers, Fig. S2c) appear mostly unchanged between tissues. Overall, these results indicate that very similar networks of specific looping interactions between CRMs within a TAD are present in cells where transcription is either active or silent.

To search for a possible explanation of these surprising results, we explored the transcription factor binding profiles of known activators and repressors in the *doc* locus ^51,61,62^. Notably, the four CRMs for which we observed looping interactions (CRM_a-d_) displayed strong binding of ‘Mothers against dpp’ (Mad) and Zerknullt (Zen), two transcriptional activators of *doc* genes that tend to localize specifically to the dorsal ectoderm at nc14 ^60,63^ (Fig. 2g). Thus, CRM hubs in the dorsal ectoderm contain *doc* activators. Therefore, contacts between *doc* promoters and CRM hubs in the dorsal ectoderm would presumably facilitate transcriptional activation (Fig. 2h). In contrast, in the mesoderm and neuroectoderm, CRM_a-d_ tend to be occupied by spatially-localized transcriptional repressors: Sna in the mesoderm, and Brinker (Brk)/Schnurri (Shn) in the neuroectoderm ^64^ (Fig. 2g) ^51,61,62^. In this context, contacts between *doc* promoters and CRM hubs in the mesoderm/neuroectoderm may instead facilitate repression (Fig. 2h).

### *Cis*-regulatory networks emerge before TADs and gene expression

Previous genome-wide and Hi-M studies have established that most *Drosophila* TADs emerge at nc14 during the major wave of zygotic gene activation (ZGA) ^38,44,45^. To explore whether the *doc*-TAD also emerges at this nuclear cycle, we performed low-resolution Hi-M experiments in embryos staged from nc11-nc12 and at nc14 (Fig. 3a), and used nuclei density to unequivocally score developmental timing (Fig. 3c, insets). Hi-M contact maps revealed that the *doc*-TAD can be detected at nc14 but not at earlier stages (Fig. 3a), thus emergence of this TAD coincides with the onset of *doc* expression (Fig. 3b).

**Figure 3.**
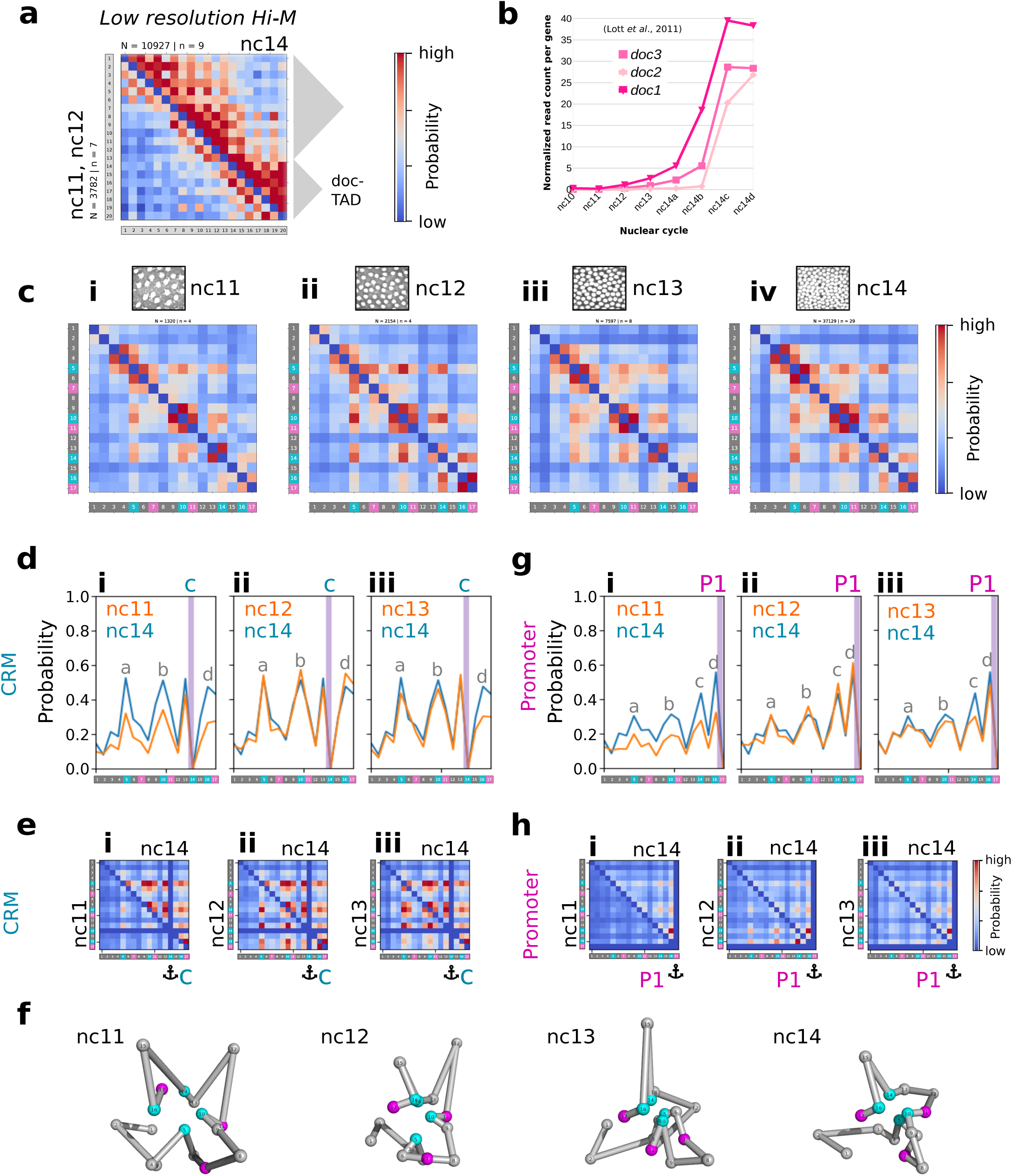
CRM loops and hubs precede TAD formation and gene expression. a. Low resolution Hi-M contact probability map of an extended genomic region around the *doc*-TAD. Upper-right map: nc14, lower-left map: nc11 and nc12. Contact probability is depicted according to the colorbar. The two TADs are depicted by gray triangles on the right. Barcodes are shown as gray stripes running on the left and bottom of the Hi-M map. N: number of cells. n: number of embryos. b. Expression profile of *doc1, doc2* and *doc3* during nuclear cycles 10-14. Nuclear cycle 14 was divided into four time-points according to the extent of cellularization (a: earliest; d: last). c. Representative images of DAPI-stained nuclei for embryos in nuclear cycles nc11 to nc14 (upper panel). High resolution Hi-M contact probability maps of the *doc*-TAD for embryos in nc11, nc12, nc13 and nc14. Barcodes are indicated below and on the left of each Hi-M map. N: number of cells. n: number of embryos. d. Comparison of 4M profiles derived from Hi-M maps at different nuclear cycles. The position of the anchor (CRM_c_) is indicated by a vertical purple line. Profiles for nc11, 12 and 13 (orange lines) are compared to nc14 (blue lines) in panels i to iii, respectively. Peaks in the profiles are annotated with the corresponding CRMs (a-d). Barcodes are indicated in the x-axis of each plot. e. Comparison of 3-way contacts between nc14 and other nuclear cycles, using CRM_c_ as anchor. Upper-right half of the matrix always depicts nc14, whereas the bottom-left half shows nc11 (panel i), nc12 (panel ii) and nc13 (panel iii). The 3-way contact probability map is color-coded according to the colorbar. The position of the anchor is indicated by a pictogram. Barcodes are indicated on the left and bottom of each map. f. Topological reconstructions of the *doc*-TAD for nc11 to nc14. CRMs and promoter regions are indicated as cyan and magenta spheres, respectively. g. Similar to panel c, anchor: *doc1* promoter (P1) h. Similar to panel d, anchor: *doc1* promoter (P1).

To determine whether specific looping interactions between CRMs appear before the emergence of TADs, we performed high-resolution Hi-M between nc11 and nc14. As our previous data showed that Hi-M maps are similar in different presumptive tissues (Fig. 2), we built Hi-M maps for the different nuclear cycles using all detectable cells independently of their location in the embryo. Surprisingly, chromatin loops between CRMs are observed very early in development (nc11) and remain almost unaffected at least until embryos reach nc14 (Fig. 3c). For example, loops between CRM_c_ and CRM_a_, CRM_b_ and CRM_d_ can be readily detected at nc11, and assume their final contact frequencies at nc12 (Fig. 3d). Similar behaviours can be observed when using other CRMs as anchors (Fig. S3a-c).

Finally, we tested whether formation of CRM hubs depends on developmental time by performing 3-way analysis using CRMs as anchors. We observed that 3-way interaction patterns are already visible at nc11, but are weaker than at nc14 (Figs. 3e, S3d-f). Indeed, patterns of 3-way interactions are almost indistinguishable from nc12 to nc14. To gather further evidence for the formation of CRM hubs during early development we obtained ShRec3D structures for each nuclear cycle. Notably, these structures show that CRMs cluster at the center of the TAD as early as nc11, with clusters becoming tighter as development progresses (Fig. 3f). Therefore, these data are consistent with clustering of CRMs forming gradually during development.

Then, we performed a similar analysis for *sna*-TAD, which also emerges at nc14 ^45^. Remarkably, pairwise contacts between *esg* and *sna* enhancers appeared at nc11 and reached their maximum intensities between nc 12-13 (Fig. S3g). As for the *doc*-TAD, we observed that the strongest 3-way contacts visible at nc14 are already present at nc11 (e.g. *sna* enhancers and the *nht* locus, white boxes, Fig. S3h), while other 3-way interactions were established at later cycles and persisted until nc14 (e.g. green boxes, Fig. S3j). In particular, 3-way interactions involving the TAD border were the last to be acquired (nc13-nc14, e.g. yellow box, Fig. S3k). All in all, these data indicate that pairwise looping interactions between CRMs in *doc* and *sna*-TADs are established from nc11 (or before) while 3-way interactions are progressively acquired. Importantly, both pairwise and multi-way looping interactions are formed before the emergence of TADs.

The timing of expression of *doc* genes coincides with the major wave of ZGA at nc14 ^65,66^. *Doc1* is the first gene to be expressed at the very beginning of nc14, while *doc3* and *doc2* are expressed towards the end of nc14, respectively (Fig. 3b). To investigate whether specific loops between *doc* promoters and CRMs displayed quantitative changes before the onset of gene expression, we plotted virtual 4M profiles with gene promoters as viewpoints. Notably, we observe that promoters contact CRMs as early as nc11, and that frequencies of interactions cease to change after nc12 (see P1 in Fig. 3g, and P2-P3 in Fig. S3a-c). 3-way interactions involving promoters could also be already detected at nc11, and become more frequent at later nuclear cycles (Figs. 3h and S3d-f). Thus, our results indicate that loops involving promoters and one or several *cis*-regulatory elements precede TAD formation and gene expression, and are equally frequent in naive pluripotent cells which do not express *doc* genes.

### Formation of CRM hubs requires the pioneer factor Zelda

Having shown that interactions between multiple CRM do not depend on transcriptional state nor developmental timing, we searched for factors that may be required for the formation of CRM hubs. The pioneer factor Zelda (Zld) has the unusual ability to overcome nucleosome barriers at specific regulatory elements, making them accessible for binding by other classical TFs prior to activation, as early as nc8-nc11 ^52–54,57,67,68^. The *doc*-TAD is particularly enriched in Zld binding, and all the CRMs displaying specific looping interactions by Hi-M appear as open chromatin and are bound by Zld (Fig. 4a). Importantly, removal of Zld leads to a decrease in *doc1-3* transcription (Fig. S4a), and to an overall reduction in chromatin accessibility, with some CRMs being particularly impacted by Zld depletion (e.g. CRM _c-d_, gray arrows in Fig. 4a) ^54,68^. To explore whether the pioneering activities of Zld were required for the establishment of CRM hubs, we performed Hi-M experiments on Zld maternally depleted embryos using *RNAi* ^67^. Given the widespread developmental defects exhibited by *Zld* RNAi embryos at stage 5 ^69^, we restricted our analysis to early nc14 *Zld RNAi* embryos.

**Figure 4.**
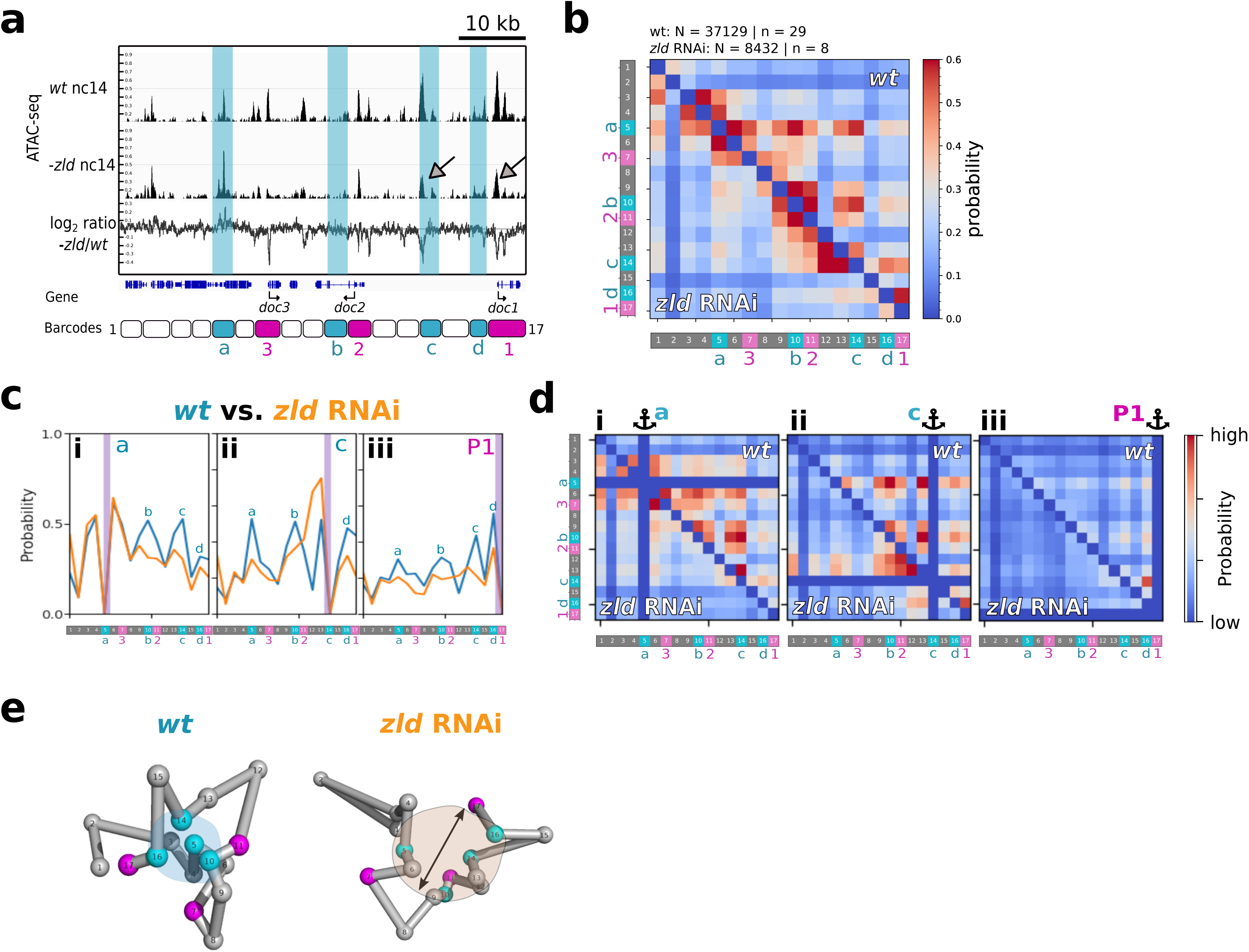
Formation of CRM loops and hubs in the *doc*-TAD requires the pioneer factor Zelda. a. ATAC-seq profiles of wildtype (*wt*, top row), *Zld* mutant (*-zld*, middle row), and log2 ratio between *-zld* and *wt* nc14 embryos (bottom row). Genomic regions occupied by barcodes containing CRM_a-d_ are annotated in cyan. The positions of the barcodes are indicated at the bottom. Two examples of peaks showing a decrease in chromatin accessibility upon Zld depletion are illustrated with gray arrows. b. High-resolution Hi-M contact probability maps for *wt* (upper-right matrix) and *zld-*RNAi (bottom-left matrix) embryos. Colormap shows contact probability. Barcodes are indicated on the left and bottom of the matrix. N: number of cells. n: number of embryos. c. 4M profiles derived from Hi-M maps for *wt* (blue) and *zld-*RNAi (orange) embryos. Anchors: CRM_a_ (panel i), CRM_c_ (panel ii) and P1 (panel iii). The position of the anchors is indicated by a vertical purple line. Barcodes are depicted on the x-axis. d. Multi-way interactions in *wt* (upper-right map) and *zld-*RNAi (bottom-left map) embryos. Anchors are indicated by a pictogram. Anchors used: CRM_a_ (panel i), CRM_c_ (panel ii) and P1 (panel iii). e. Topological reconstructions of *doc*-TAD derived from Hi-M matrices from *wt* and *zld-*RNAi embryos (panel b). CRMs and promoter regions are indicated by cyan and magenta spheres, respectively. The CRM hub is indicated by a salmon shading in the *wt* reconstruction. The arrow indicates a separation of CRMs in the absence of Zld.

Strikingly, we observed large changes in the Hi-M contact matrix, with most specific looping interactions between CRMs disappearing upon depletion of Zld (Fig. 4b). To quantify this effect, we performed 4M analysis using promoters and CRMs as viewpoints. In all cases, we observed a reduction in the interaction frequency between promoters and CRMs upon Zld depletion (Figs. 4c-iii, S4b-i, ii, iii). Interestingly, interactions between CRMs were also disrupted to a large degree (Figs. 4b-c, S4b-v,vi,vii,viii), with the only peaks remaining corresponding to the first neighbours from the anchor, most likely reflecting the polymer nature of the chromatin fiber and not specific looping interactions. For instance, upon Zld depletion CRM_c_ showed the largest drop in ATAC-seq signal amongst CRMs (Fig. 4a), and its interactions with other CRMs appear almost absent in Zld-RNAi embryos (Figs. 4b, 4c-ii). Similarly, CRM_a_ exhibited also a large decrease in interaction frequencies with other CRMs (Fig. 4c-i), despite a mild increase in ATAC-seq signal upon Zld depletion (Fig. 4a). Finally, formation of CRM hubs was also considerably impacted in Zld-RNAi embryos (Figs. 4d and S4c), with topological reconstructions showing a loss of CRM clustering upon depletion of Zld (Fig. 4e). All together, these results suggest a model whereby the pioneering activity of Zld participates in the formation of CRM loops and hubs during early embryogenesis, possibly through its ability to open chromatin at specific CRMs.

## Discussion

In this study, we use a novel high-resolution, imaging-based, single-cell spatial genomics approach (Hi-M) to link chromosome topology and transcriptional regulation during early *Drosophila* development. We reveal extensive interaction networks within developmental TADs primarily involving *cis*-regulatory modules. Critically, these networks arise thanks to the spatial clustering of multiple enhancers (CRM hubs) and are mostly invariant during cell fate specification and gene activation. Networks of pairwise CRM contacts and CRM hubs arise during early development, before the onset of gene expression and before the emergence of TADs, and require the pioneering activity of the transcription factor Zld.

One of the important surprises of this study is that physical proximity between multiple CRMs and promoters is observed with very similar frequencies in cells with three distinct fates. This network of contacts appeared during early embryogenesis and within the regulatory unit defined by single TADs, which tend to be small in *Drosophila* (∼70 kb) ^38,70,71^. These results are consistent with those obtained at later stages of *Drosophila* embryogenesis, showing that enhancers located at considerably larger distances (∼100 kb) can also form binary loops that are present in cells from different tissues ^17^. Similarly, E-P interactions at the mouse *HoxD* locus were detected in tissues where target genes were not expressed ^18^. From a developmental perspective, the formation of loops between promoters and distal regulatory elements in cells where genes need to be repressed can be seen as a ‘dangerous liaison’. Indeed, once a loop is established, transcriptional activation could rapidly occur in cells where that specific promoter should be kept inactive.

This apparent dichotomy, however, can be rationalized in terms of the spatio-temporal patterning of the *cis*-regulatory logic of transcription factors during embryogenesis. For instance, in mesodermal cells, most *doc* CRMs are bound by the spatially-localized transcriptional repressor Sna ^42^, which acts as a silencer in the mesoderm. In this case, communication between promoters and distal CRMs may reinforce transcriptional repression rather than enhancing transcriptional activation. This interpretation is in agreement with the recent finding that the vast majority of enhancers can act as silencers in alternate cell types during *Drosophila* development ^72^, however other silencing mechanisms may also be at play ^73^. Thus, we hypothesize that the optimal mechanism to ensure both rapid and efficient activation and repression during development may involve two steps: the rapid priming of key CRMs via ubiquitously maternally-deposited pioneer factors (e.g. Zld), followed by regulation of transcriptional output by spatially- and temporally-localized transcriptional activators and repressors. In this model, 3D chromatin architecture plays a double role as 3D contacts would serve to simultaneously reinforce both activation and repression at a particular developmental stage while allowing for flexibility at later stages. For instance, a repressive CRM loop in a tissue at an early developmental stage may switch to a CRM loop with activation capacities at later stages by changing transcription factor occupancy. Future experiments testing whether CRM loops and hubs display more differences in active and repressed tissues at later stages of development will be important to validate these hypotheses.

Previous studies suggested that invariant E-P loops may be pre-established and stable ^14,17,74,75^. In agreement with these results, our data indicate that E-P loops can form early, well before the onset of gene expression. However, in all cases, we measured low frequencies of looping interactions between functional elements. These results are consistent with previous measurements of absolute contact frequencies within TADs and between E-P ^47,76–78^. Thus, these results indicate that different sets of multi-way E-E and E-P contacts are present in different cells, and that these contacts may be highly dynamic.

Recent studies reported the existence of enhancer hubs: spatially-localized clusters containing multiple enhancers ^27,28,31^ that may facilitate transcriptional activation by creating a local microenvironment whereby transcriptional resources are shared, akin to early models of ‘transcription hubs’ ^79^. Formation of enhancer hubs may require interactions between components of the transcriptional machinery which could contribute, or result from, the assembly of phase-separated condensates ^30,35,36,80–82^. In this model, enhancers need not directly touch their target promoters but merely come into close proximity ^24,83^. Overall, these findings and models are consistent with our observation that multiple endogenous CRMs within a TAD come together in space to form hubs in single, actively-transcribing cells. Most notably, we also observed the formation of similar hubs in inactive cells, suggesting that repressive elements may also form spatially-localized clusters of transcriptional repressors to share resources and reinforce their silencing activities.

In *Drosophila*, TADs emerge concomitantly with the major wave of zygotic gene activation (ZGA) ^38,44,45^. Previous studies reported the existence of chromatin loops typically at considerably large genomic distances spanning two or more TADs ^17,44^ or concerning Polycomb binding sites ^44,84^. Here, we observed that chromatin loops between *cis*-regulatory elements within *Drosophila* TADs are widespread, mimicking the common CTCF-mediated chromatin loops present within mammalian TADs ^15,39^. In addition, we found that multiple CRMs can cluster together to form *cis*-regulatory hubs located at the interior of TADs, suggesting a mechanism to sequester enhancers in space to reduce the activation of genes in neighboring TADs. Importantly, formation of CRM hubs precedes the emergence of TADs, consistent with the finding in mammalian cells that subsets of E-P contacts arise rapidly after mitosis before TADs are reformed ^85^. Thus, our results suggest that CRM hubs and TADs likely form by different mechanisms. All in all, we hypothesize that CRM hubs represent an additional functional level of genome organization independent of TADs. This additional layer can also be regulated by priming of enhancers and promoters by paused polymerases ^86–88^ or pioneer factors ^52,53^, as well as by chromatin marks ^89^.

Interestingly, we observed that CRM hubs, as well as interaction networks between CRMs and cognate promoters are established very early in naive pluripotent nuclei, prior to cell fate commitments. Critically, preferential loops and CRM hubs were considerably attenuated upon depletion of the pioneer factor Zld. We and others have recently shown that Zld forms nuclear hubs in early *Drosophila* embryos ^33,34^, and that Zld hubs are re-established by the end of mitosis, prior to any sign of transcriptional activation. Taken together, our results suggest a model whereby Zld could foster the formation of CRM hubs by rendering chromatin accessible during early development, as a first step of cell specification to ensure maximum plasticity. Future work involving the detection of a larger number of CRMs will be needed to elucidate the factors and mechanisms involved in spatial clustering of developmental CRMs into nuclear micro-environments.

This study provides clear evidence of the advantages of imaging-based technologies (e.g. Hi-M) to detect multi-way chromatin loops and transcriptional output with spatial resolution. First, this technique enables the detection of interactions in cells within different tissues and in distinct transcriptional states in a single experiment. Second, this tool may serve to discover novel *cis*-regulatory modules, and to assess whether predicted CRMs actually contact their target promoters and measure its activity in a given tissue and at a specific developmental time, without the burden of genetic manipulation. Third, the ability to detect multi-way interactions will be important to further dissect the mechanisms of transcriptional control by distal CRMs, as well as mechanisms of transcriptional co-regulation. Finally, combination of these approaches with opto-genetic manipulation will open exciting avenues for the spatio-temporal control of regulators to assess their roles in shaping 3D architecture and regulating transcription.

## Methods

### *Drosophila* stocks and embryo collection

Fly stocks were maintained at room temperature (RT) with natural light/dark cycle and raised in standard cornmeal yeast medium. The *yw* stock was used as a control. Zelda depleted embryos were obtained from females from the cross between *nos-Gal4*:VP16 (BL4937) and *UASp-shRNA-zld* ^67^. After a pre-laying step, flies were allowed to lay eggs for 1.5 h on new yeasted 0.4% acetic acid plates. Embryos were then incubated at 25 °C until they reached the desired developmental stage. Embryos were collected and fixed as previously described ^46^. Briefly, embryos were dechorionated with bleach, rinsed and fixed with a 1:1 mixture of 4% methanol-free formaldehyde in PBS and Heptane. Embryos were stored in methanol at −20 °C until further use.

### Hi-M libraries

Oligopaint libraries, consisting of unique ∼35/41-mer sequences with genome homology, were obtained from the Oligopaint public database (http://genetics.med.harvard.edu/oligopaints). We selected 20 barcodes in the *doc locus* (3L: 8882600..9039000 *Drosophila* release 5 reference genome in all cases) for the low-resolution Hi-M library, 17 barcodes encompassing the *doc*-TAD (3L: 8974562..9038920) for the high-resolution Hi-M library, and 65 barcodes (2L:15244500..15630000) for the high-resolution *sna* locus library. For each barcode, we used 45-50 probes, covering ∼3 kb. One of the barcodes in the low-res library was selected as the fiducial barcode to use for drift correction (see below), whereas an additional fiducial barcode located ∼1 Mb away was used in the high-res libraries for drift correction. The coordinates of the targeted genomic regions are listed in Supplementary Table 2.

Each oligo in the pool consisted of 5 regions: i-a 21-mer forward priming region, ii-a 32-mer (low-res library) or two 20-mer separated by an AT sequence (high-res libraries) readout region unique for each barcode, iii-a 35/41-mer genome homology region, iv-a 32-mer (low-res library) or 20-mer (high-res libraries) readout region and v-a 21-mer reverse priming region. The designed template oligo pools were ordered from CustomArray. The procedure to amplify oligo pools to obtain the primary libraries was as previously described ^46^. It involved a 4-step procedure consisting of i-limited-cycle PCR, ii-amplification via T7 *in vitro* transcription, iii-reverse transcription and iv-alkaline hydrolysis and purification. The sequence of the primers used for amplification of the libraries are listed in Supplementary Table 3.

For the low-resolution library, we employed 21 unique Alexa647-labeled sequence oligos (imaging oligos), complementary to the readout region present in the primary oligo. The fluorophore was attached via a disulfide linkage cleavable by the mild reducing agent Tris(2-carboxyethyl)phosphine (TCEP), using a previously described strategy ^46^. Alternatively, for the high-resolution libraries, we used “adapter” oligos, consisting in a 20-mer region complementary to the readout sequence able to recognize the barcode being targeted, a 10-mer spacer sequence and a 32-mer region able to bind to a unique Alexa647-labeled oligo (containing a disulfide linkage). In this approach, a single fluorescent oligo is required ^47^. For fiducial barcodes, a non-cleavable Rhodamine-labeled oligo was used. The sequences of the imaging and adapter oligos are listed in Supplementary Table 4. PCR and reverse transcription primers primers used in probe synthesis, as well as adapter oligos and fluorescently-labeled oligos, were purchased from Integrated DNA Technology (IDT). The whole set of Oligopaints used can be found in Supplementary Table 5.

### RNA-FISH probes

RNA probes were obtained by *in vitro* transcription from a vector containing the sequences targeting *sna* (previously described in^45^, *doc1, doc2* or *doc3* genes in the presence of digoxigenin (DIG) or biotin (BIO) haptenes. Vector was linearized before the *in vitro* transcription with a specific restriction enzyme. RNA probes produced in this manner were then treated with carbonate buffer at 65 °C for 5 min (*sna* probe) or for 2 min (*doc1,doc2,doc3* probes). The information on each probe, including the primers used to clone the target sequences, are listed in Supplementary Table 6.

### RNA Fluorescent *In situ* Hybridization

*In situ* hybridization was as described previously ^46^, with modifications to allow for the detection of two different species of RNA. The reader is invited to read our detailed protocol in the aforementioned reference. Briefly, fixed embryos were passed through 1:1 mixture of methanol:ethanol and then pure ethanol. Embryos were then post-fixed with 5% formaldehyde in PBT (PBT = 0.1% Tween-20 PBS) for 25 min. Then, embryos were incubated 4 times with PBT during 15 min and permeabilized 1 h with 0.3% Triton in PBS. Embryos were rinsed with PBT and incubated for 2 h with RHS at 55 °C (RHS = 50% formamide, 2X SSC, 0.1% Tween-20, 0.05 mg/ml heparin, 0.1 mg/ml salmon sperm). In the meanwhile, RNA probes were heated at 85 °C for 2 min, transferred to ice for 2 min and then incubated with the embryos in RHS for 16-20 h at 55 °C for RNA hybridization. The next day, embryos were washed 4 times with RHS at 55 °C and 3 times with PBT at RT. Then, a saturation step was performed with blocking solution (blocking reagent Sigma #11096176001, 100 mM Maleic acid, 150 mM NaCl, pH = 7.5) for 45 min.

Then the protocol depends on whether embryos were used for Hi-M (*sna*/*doc1* double labeling) or to reveal *doc1, doc2* and *doc3* expression patterns (Fig. S1b). To reveal the expression patterns of *doc* genes, the combination of *doc1*-DIG/d*oc2*-BIO or *doc2*-DIG/*doc3*-BIO was used. After the saturation step, embryos were incubated with primary antibodies at 1:375 dilution (sheep anti-DIG, Roche cat #11333089001 and mouse anti-Biotin, Life technologies cat #03–3700) overnight at 4°C. The next day embryos were washed 6 times in PBT for 10 min. Embryos were incubated 1 h in blocking solution, then 2 h with secondary antibodies at 1:500 dilution (anti-mouse Alexa488-conjugated Life technologies cat #A21202 and anti-sheep Alexa555-conjugated Life technologies cat #A21436) and washed 6 times in PBT. Finally, embryos were incubated 10 min with a 0.5 mg/mL DAPI solution, washed with PBT and mounted in ProLong(tm) Diamond Antifade.

For Hi-M, both *sna* and *doc1* probes were DIG-labeled. By taking advantage of the differential spatial expression pattern, we labeled both RNAs simultaneously by the combination of both probes during incubation and the use of a single anti-DIG antibody and a tyramide signal amplification (TSA) reaction. After RNA hybridization and the saturation step, the activity of endogenous peroxidases was eliminated by incubating with 1% H_2_O_2_ in PBT for 30 min. After rinsing with PBT, embryos were incubated overnight at 4 °C with sheep anti-DIG conjugated with POD (Sigma-Aldrich cat #11207733910) with 1:500 working dilution in PBT. The next day, embryos were washed with PBT and incubated for 30 min with tyramide-coupled Alexa 488. Next, H_2_O_2_ was added to a final concentration of 0.012% during another 30 min. Embryos were washed with PBT and stored at 4 °C until further use.

### Hybridization of Hi-M primary library

Hybridization followed a previously described protocol ^46^. Briefly, embryos were RNase treated for 2 h, permeabilized 1 h with 0.5% Triton in PBS and rinsed with sequential dilutions of Triton/pHM buffer to 100% pHM (pHM = 2X SSC, NaH_2_PO_4_ 0.1 M pH = 7, 0.1% Tween-20, 50% formamide (v/v)). Embryos in pHM were preheated at 80 °C, the supernatant was aspirated and 30 *μ*L of FHB (FHB =50% formamide, 10% dextran sulfate, 2X SSC, salmon sperm DNA 0.5 mg/mL) containing 225 pmol of the primary library was pipetted directly onto the embryos. Mineral oil was added on top and the tube was incubated overnight at 37 °C. The next day, oil was carefully removed and embryos were washed two times during 20 min at 37 °C with 50% formamide, 2X SSC, 0.3% CHAPS. Next, embryos were sequentially washed for 20 min at 37 °C with serial dilutions of formamide/PBT to 100% PBT. Embryos were rinsed with PBT and stored at 4 °C until the imaging step.

### Imaging system

All experiments were performed on a home-made wide-field epifluorescence microscope built on a RAMM modular microscope system (Applied Scientific Instrumentation) coupled to a microfluidic device as described previously ^45,46^. Samples were imaged using a 60x Plan-Achromat water-immersion objective (NA = 1.2, Nikon, Japan). The objective lens was mounted on a closed-loop piezoelectric stage (Nano-F100, Mad City Labs Inc. - USA). Illumination was provided by 4 lasers (OBIS-405/488/640 nm and Sapphire-LP-561 nm, Coherent – USA). Images were acquired using a sCMOS camera (ORCA Flash 4.0V3, Hamamatsu – Japan), with a final pixel size calibrated to 106 nm. A custom-built autofocus system was used to correct for axial drift in real-time and maintain the sample in focus as previously described ^45^.

A fluidic system was used for automated sequential hybridizations, by computer-controlling a combination of three eight-way valves (HVXM 8-5, Hamilton) and a negative pressure pump (MFCS-EZ, Fluigent) to deliver buffers and secondary readout probes onto a FCS2 flow chamber (Bioptechs). Software-controlled microscope components, including camera, stages, lasers, pump, and valves were run using a custom-made software package developed in LabView 2015 (National Instrument).

### Acquisition of Hi-M datasets

Embryos were attached to a poly-L-lysine coated coverslip and mounted into the FCS2 flow chamber. Fiducial readout probe (25 nM Rhodamine-labeled probe, 2X SSC, 40% v/v formamide) was flowed onto the sample and hybridized for 15 min, washed for 10 min with readout washing buffer (2X SSC, 40% v/v formamide) and for 5 min with 2X SSC before injecting 0.5 mg/mL DAPI in PBS to stain nuclei. The imaging buffer (1x PBS, 5% w/v glucose, 0.5 mg/mL glucose oxidase and 0.05 mg/mL catalase) was injected. Subsequently, 10-15 embryos were selected according to developmental stage and orientation and segmented into a mosaic of multiple fields of view (FOV of 200 x 200 *μ*m). After bright field image recording, z-stacks were taken with 405, 488 and 561 nm laser illuminations. The z-stacks had a step size of 250 nm with a total range of 15 μm.

Next, the sample was sequentially hybridized with different secondary readout probes, imaged in the Rhodamine and the Alexa-647 channels, and photobleached. For each round of secondary hybridization, the sample was treated with secondary hybridization buffer (25-50 nM imaging oligo, 2X SSC, 40% v/v formamide, that also included 50 nM of adapter oligo in the case of the high-res libraries, see *Hi-M libraries*,) for 15 min, then washed with readout washing buffer and with 2X SSC before injecting imaging buffer. After imaging, the fluorescence of the readout probes was extinguished using a chemical bleaching buffer (2X SCC, 50 mM TCEP hydrochloride) for 10 min and then the sample was washed with 2X SSC for 5 min before a new hybridization cycle started. All buffers were freshly prepared and filtered for each experiment. The imaging buffer used for a single experiment was stored under a layer of mineral oil and renewed every 12-15 h. Further details can be found on our previously published protocol ^46^.

### Image processing

Our home-made Hi-M microscope produced z-stacks in DCIMG format, which were converted to TIFF using proprietary software from Hamamatsu. TIFF images were then deconvolved using Huygens Professional version 20.04 (Scientific Volume Imaging, the Netherlands, https://svi.nl/). Further analysis steps were performed using a homemade analysis software that implemented the steps described previously ^46^. Briefly, images were first z-projected using either sum (DAPI channel) or maximum intensity projection (barcodes, fiducials). Image-based cross-correlation was used to align the fiducial channels. These corrections were then used to align DAPI and barcode images. Next, the positions of the XY centers of barcodes were detected with sub-pixel resolution using local maximum fitting functions from the ASTROPY package ^90^. Nuclei were segmented from projected DAPI images by adaptive local thresholding and watershed filtering ^46^. RNA images were segmented by manually drawing polygons over the nuclei displaying a pattern of active transcription. Barcodes and RNA status were then attributed to each single nuclei by using the XY coordinates of the barcodes, the projected DAPI masks of nuclei, and the transcriptional status from manual masking. Finally, pairwise distance matrices were calculated for each single cell and converted into contact frequency maps by using a threshold of 250 nm. Contact frequencies obtained using this pipeline and those using previous pipelines ^46^ produced highly correlated results. Image processing was carried out from Linux terminals connected to a server running Linux PopOS 19.10, with 2-4 GeForce GTX 1080Ti GPU cards. Statistical evaluation of Hi-M datasets was performed using a bootstrapping approach (Fig. S1h).

### 4M profiles and multi-way interactions

4M profiles were obtained by slicing the corresponding Hi-M contact map across a given anchor. Multi-way interactions were obtained by selecting an anchoring barcode and calculating the single-cell pairwise distances to all possible pairs of barcodes. If both barcode-anchor distances for a given barcode pair in a single cell are below the contact threshold (250 nm), this cell is considered to have a 3-way interaction for this anchor and barcode pair. The 3-way contact frequency is then obtained by dividing the number of cells that show a 3-way interaction by the number of cells where the three barcodes involved in the 3-way interaction have been detected.

### ShRec3D Structures

Three-dimensional topological representations were obtained from Hi-M pairwise distance maps using our own Python implementation of the approaches described by Lesne *et al*. and Morlot *et al*. for ShRec3D ^59,91^. Starting from the single-cell pairwise distance matrix, an ensemble of pairwise distance matrices were calculated using the first maximum of the kernel density estimation. These pairwise distances were converted into 3D coordinates for each barcode using nonclassical metric multidimensional scaling. When necessary, structures were mirrored and a ball-and-stick representation was rendered with PyMOL (The PyMOL Molecular Graphics System, Version 2.3 Schrödinger, LLC.).

## Acknowledgements

We are grateful to N. Benabdallah, G. Cavalli, T. Forne, T. Robert, J. Bonnet and members of the Lagha and Nollmann labs for their critical reading of the manuscript. This project was funded by the European Union’s Horizon 2020 Research and Innovation Program (Grant ID 724429) (M.N.). We acknowledge the Bettencourt-Schueller Foundation for their prize ‘Coup d’élan pour la recherche Française’, the France-BioImaging infrastructure supported by the French National Research Agency (grant ID ANR-10-INBS-04, ‘‘Investments for the Future’’), and the *Drosophila* facility (BioCampus Montpellier, CNRS, INSERM, Univ Montpellier, Montpellier, France). M.G. was funded by the Deutsche Forschungsgemeinschaft (DFG, German Research Foundation) - project ID 431471305. ML’s lab is supported by an ERC_Starting Grant (SyncDev) and CNRS. M.B. is supported by an FRM fellowship. A.M.C.G. is a postdoctoral fellow of Consejo Nacional de Investigaciones Científicas y Técnicas (CONICET), Argentina.

## Author Contributions

Conception and design: A.M.C.G., M.L., M.N.; Acquisition of data: S.M.E., C.H., M.B.; Analysis: M.G., S.M.E., M.N.; Software: M.G., M.N., J.B.F.; Interpretation of data: S.M.E., M.G., M.N., M.L.; Writing: M.L. and M.N.; Visualization: S.M.E., M.G., M.N.; Supervision: M.N. and M.L; Fund acquisition: M.N and M.L.

