## Supplementary Data for "*Cis*-regulatory chromatin loops arise before TADs and gene activation, and are independent of cell fate during development"

### SUPPLEMENTARY INFORMATION RELATED TO

### Cis-regulatory interaction networks are independent of cell fate during early embryogenesis, and arise before TADs and gene activation

Sergio Martin Espinola[1*], Markus Götz[1*], Jean-Bernard Fiche[1], Maelle Bellec[2], Christophe Houbron[1], Andrés M. Cardozo Gizzi[3], Mounia Lagha[2#], Marcelo Nollmann[1#]

** Co-first authors*

#### SUPPLEMENTARY FIGURE 1

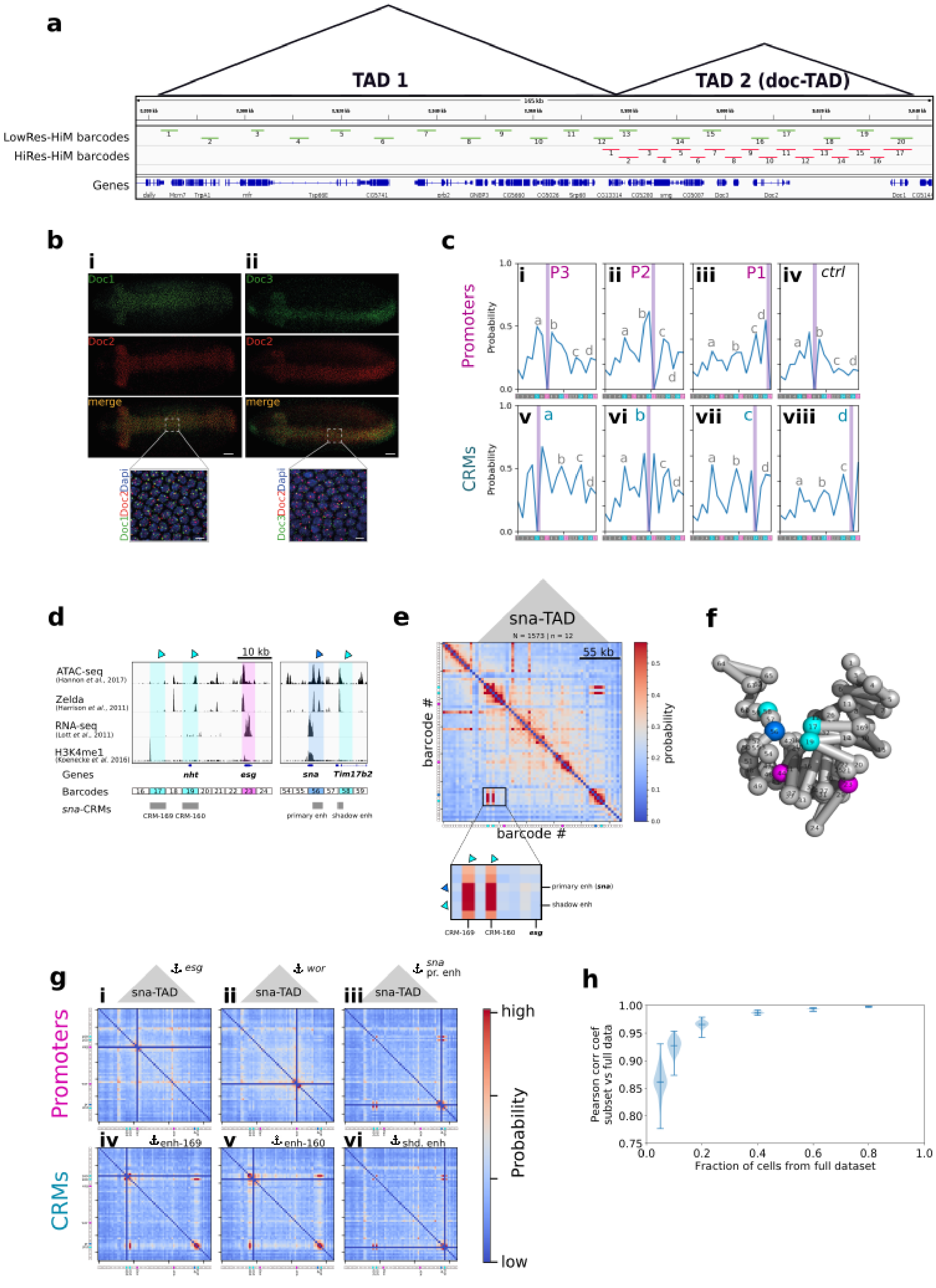


**a,** Schematic representation of the genomic positions of barcodes for the low (green) and high (red) resolution *doc* Hi-M libraries. Triangles demarcate the two TADs registered in this genomic region [1]. Genes are shown below the barcodes. **b,** RNA expression patterns of *doc1*, *doc2* and *doc3* in late nc14 wild-type embryos. Double RNA FISH staining (panel i:

*doc1* and *doc2*; panel ii: *doc2* and *doc3*), scale bar 50 $\mu$m. Inset, merge of *doc1*, *doc2* and DAPI signals (panel i) or *doc3*, *doc2* and DAPI signal (panel ii), scale bar 5 $\mu$m. Images were taken on a Zeiss LSM880 confocal. **c,** 4M profiles derived from Hi-M maps of dorsal ectoderm cells (expressing *doc1*) in nc14. Anchors were placed at P3 (panel i), P2 (panel ii), P1 (panel iii), control barcode (panel iv), $CRM_a$ (panel v), $CRM_b$ (panel vi), $CRM_c$ (panel vii), $CRM_d$ (panel viii). Anchors are indicated by vertical purple lines. **d,** Epigenetic profile of selected regions around the *esg* and *sna* genes within the *sna*-TAD. Accessibility (ATAC-seq), pioneer factor binding (Zelda), transcriptional activity (RNA-seq), chromatin marks for active enhancers marks (H3K4me1), and for the transcriptional activators Dorsal, Zen and Mad are shown. A subset of barcodes were annotated as cis-regulatory modules (shown in cyan): **$CRM_{169}$** harbors the canonical H3K4me1 active enhancer mark; **$CRM_{160}$**, and **shadow** *sna* enhancer were described in the RedFly database. One barcode (magenta) was assigned as harbouring the *esg* promoter. One barcode (blue) was assigned as having both the *sna* promoter and its primary enhancer. See Supplementary Table 1 for more details. **e,** Hi-M contact probability map of the *sna* locus color coded according to the scale bar on the right. Inset shows a magnified view of a region containing CRMs around *esg* and *sna*. N: number of cells, n: number of embryos. **f,** 3D topological representation of the *sna*-TAD. Bead colors are as in panel d. Barcode 44 contains the *wor* promoter. **g,** Multi-way interactions between promoters (panels (i-iii) and CRMs (panels iv-vi). Anchoring barcodes are highlighted by a pictogram and a blue cross on the map. Barcodes are indicated on the left and bottom axes. **h**, Pearson correlation coefficient of the contact probability of the full *doc*-TAD hi-resolution Hi-M dataset (nc14, dorsal cells displaying *doc1* expression) against subsets with a fraction of the cells taken from the whole dataset. One hundred random subsets were generated for each tested subset size. The error bars indicate the mean and extreme values of the distribution.

#### SUPPLEMENTARY FIGURE 2

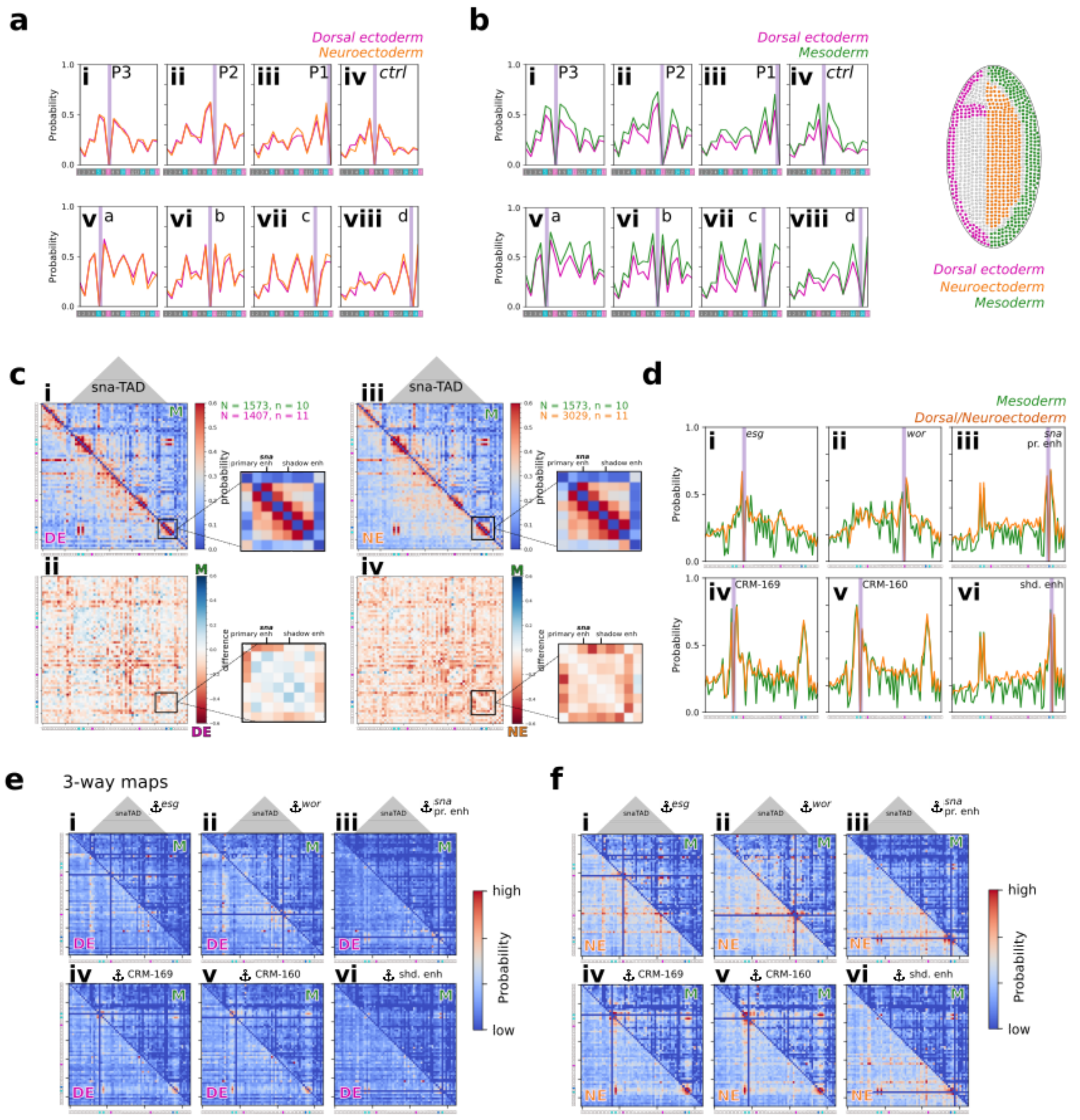


**a,** Comparison of 4M profiles between DE (magenta) and NE (orange) for different anchors within the *doc*-TAD (Panels i-iii: promoters. Panel iv: control. Panels v-viii: CRMs). **b,** Comparison of 4M profiles between DE (magenta) and M (green) for different anchors within the *doc*-TAD (Panels i-iii: promoters. Panel iv: control. Panels v-viii: CRMs). Right panel: scheme indicating the three presumptive tissues. **c,** Panel i: Hi-M contact probability map of the *sna* locus for M (upper-right map) versus DE (lower-left map). Inset show a magnification of the region around *sna*. Panel ii: Same but for the difference between M and DE Hi-M maps. Blue indicates larger contact probabilities in M whereas red indicates larger contact probabilities in DE. Panel iii: Similar to panel i, but for M (upper-right map) versus NE (lower-left map). Panel iv: Similar to panel ii, but for M versus NE. **d,** Comparison of 4M profiles between M (green) and DE/NE(orange). Anchors within the *sna*-TAD are indicated in each panel by vertical purple lines. A subset of peaks is annotated using the nomenclature from Figure 1d. **e,** Comparisons of multi-way maps for M (upper-right map) versus DE (lower-left map) in the *sna* locus using the anchors indicated in each panel by pictograms and dark blue crosses. Maps are color-coded according to the scale bar on the right. **f,** Similar to panel e, but for M (upper-right map) versus NE (lower-left map).

#### SUPPLEMENTARY FIGURE 3

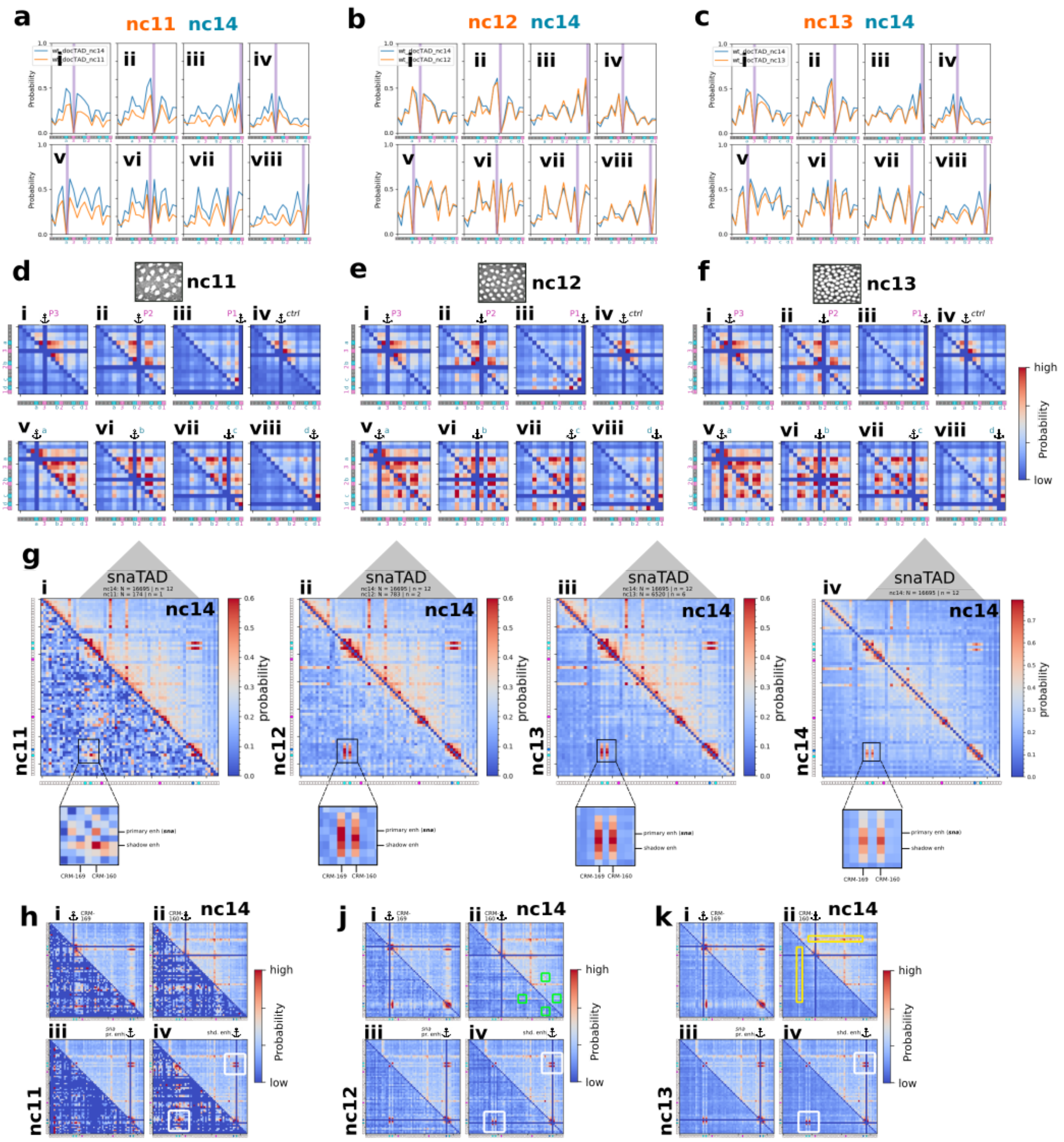


**a,** Comparison of 4M profiles between embryos in nc14 (blue lines) and nc11 (orange) for different anchors within the *doc*-TAD (Panels i-iii: promoters. Panel iv: control. Panels v-viii: CRMs). The position of the anchor is indicated by a vertical purple line. **b,** Similar to panel A, but comparing 4M profiles between embryos in nc14 (blue lines) and nc12 (orange). **c,** Similar to panel A, but comparing 4M profiles between embryos in nc14 (blue lines) and nc13 (orange). **d,** Comparison of multi-way interaction matrices of nc14 (upper-right map) and nc11 (lower-left map). Anchors (dark blue crosses) are as follows: *doc3, doc3, doc1* promoters (panels i-iii), control region (panel iv), $CRM_{a\text{-}d}$ (panels v to viii). Representative image of DAPI-stained nuclei for nc11 is shown on top. Barcodes are shown on the left and bottom of multi-way maps. **e,** Similar to panel d, but for nc14 (upper-right map) and nc12 (lower-left map). Representative image of DAPI-stained nuclei for nc12 is shown on top. **f,** Similar to panel d, but for nc14 (upper-right map) and nc13 (lower-left map). Representative image of DAPI-stained nuclei for nc13 is shown on top. **g,** Comparison of Hi-M contact probability maps in the *sna* locus for nc14 (upper-right map) and nc11 (panel i), nc12 (panel ii), nc13 (panel iii) and 14 (panel iv) (lower-left maps). Maps are color-coded according to the scale bar on the right. Inset on the bottom of each map shows a magnification of the region around *esg* and *sna* CRMs (see Supplementary Figure 1 and Supplementary Table 1). **h,** Comparison of multi-way contact maps between nc14 (upper-right maps) and nc11 (lower-left maps). Maps are color-coded according to the scale bar on the right. The position of anchors are indicated by dark blue crosses. White boxes indicate contacts already present at nc11 that persist through nc14. **j,** Similar to panel h, but comparing nc14 to nc12. Green boxes indicate contacts that emerge at nc12 and persist at nc14. **k,** Similar to panel h, but comparing nc14 to nc13. Yellow boxes indicate interactions that appear at nc13 (at the TAD border).

**SUPPLEMENTARY FIGURE 4**

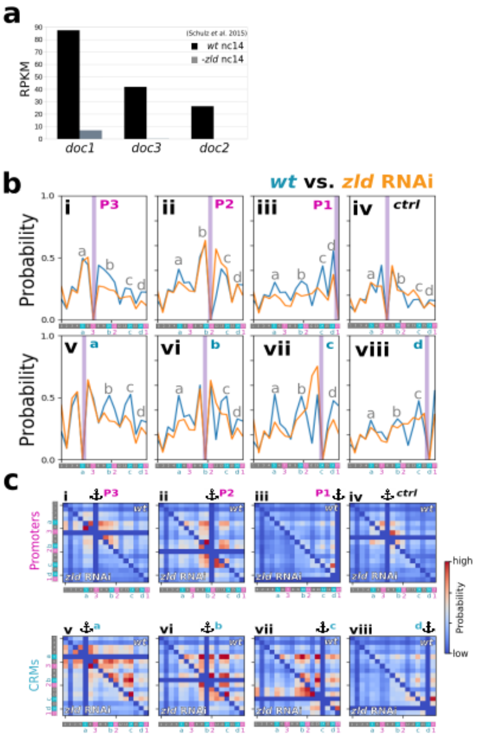


**a,** Transcription levels (RNAseq) of *doc1, doc2* and *doc3* in wild-type versus -*zld* embryos [2]. **b,** Comparison between 4M profiles of wild-type embryos (wt, blue) and of *zld* RNAi embryos (orange). The position of anchors within the *doc*-TAD are indicated by vertical purple lines. As in previous figures, promoters are represented in panels i-iii, a control locus in panel iv, and CRMs in panels v to viii. **c,** Comparison between multi-way interaction maps of wild-type embryos (upper-right maps) and *zld* RNAi embryos (lower-left maps). Anchors are indicated by pictograms and dark blue crosses. As in previous figures, promoters are represented in panels i-iii, a control locus in panel iv, and CRMs in panels v to viii.

**SUPPLEMENTARY TABLE 1. Cis-regulatory modules in *doc-* and *sna*-TADs.**

The list of enhancers was obtained from the REDFly database [3]. Enhancers highlighted in blue correspond to the HiRes-HiM barcodes 10 ($CRM_b$) 14 ($CRM_c$), and 16 ($CRM_d$) for *doc*-TAD; and 19 ($CRM_{160}$), 56 (primary), and 58 (shadow) for *sna*-TAD. No registers were found for barcode 29 ($CRM_a$) in *doc*-TAD, and barcode 17 ($CRM_{169}$) in *sna*-TAD.

| Enhancer name (*doc-* TAD) | Coordinates (dm3) | | Name in this study |
|---|---|---|---|
| | Start | End | |
| Doc3_Doc3_At16 | 8996290 | 8997200 | |
| Doc3_7731 | 8996498 | 8997275 | |
| Doc2_ae7 | 9008069 | 9011039 | ***$CRM_b$*** |
| Doc2_Cluster-8519 | 9009498 | 9010463 | |
| Doc2_7739 | 9009906 | 9010650 | |
| Unspecified_VT27864 | 9013475 | 9016012 | |
| Unspecified_GMR44H04 | 9014508 | 9017818 | |
| Unspecified_VT27865 | 9015358 | 9017515 | |
| Unspecified_GMR44G06 | 9016846 | 9019977 | |
| Unspecified_VT27867 | 9018687 | 9020860 | |
| Unspecified_GMR44B11 | 9019444 | 9021971 | |
| Doc1_GMR46B02 | 9021027 | 9024903 | ***$CRM_c$*** |
| Doc1_Doc_DocF4s1 | 9022917 | 9023482 | |
| Doc1_GMR45A12 | 9023477 | 9026879 | |
| Unspecified_VT27870 | 9023654 | 9025855 | |
| Doc1_7750 | 9023860 | 9024486 | |
| Doc2_Van53 | 9025299 | 9027490 | |
| Doc2_ae8 | 9025373 | 9027410 | |
| Unspecified_VT27871 | 9025463 | 9027668 | |
| Doc1_GMR43H05 | 9026063 | 9028354 | |
| Doc1_GMR44D03 | 9026926 | 9029637 | |
| Unspecified_VT27872 | 9027116 | 9029229 | |
| Unspecified_VT27873 | 9028822 | 9030936 | ***$CRM_d$*** |
| Doc1_GMR45H05 | 9029082 | 9032871 | |
| Doc2_ae9 | 9030154 | 9032191 | |
| Doc1_Doc1-1-lacZ | 9031836 | 9032635 | |
| Doc1_7753 | 9032001 | 9032591 | |
| Doc1_GMR43G08 | 9032007 | 9034177 | |
| Unspecified_VT27876 | 9034413 | 9037584 | |
| Doc1_GMR43B04 | 9034524 | 9035168 | |
| Doc1_GMR43E04 | 9035179 | 9036749 | |
| Doc1_Doc1-2-lacZ | 9035275 | 9036074 | |
| Doc2_ae10 | 9035288 | 9036483 | |
| Unspecified_Van54 | 9035320 | 9036469 | |
| Doc1_GMR43A11 | 9036901 | 9037501 | |

| Enhancer name (*sna*-TAD) | Coordinates (dm3) | | Name in this study |
|---|---|---|---|
| | Start | End | |
| nht_120bp | 15318847 | 15318966 | **$CRM_{160}$** |
| Unspecified_VT7839 | 15318861 | 15321392 | |
| esg_enhancerG | 15319131 | 15321427 | |
| Unspecified_VT7840 | 15320804 | 15322907 | |
| esg_D7 | 15320871 | 15323006 | |
| esg_D5 | 15320871 | 15324419 | |
| esg_D1 | 15320871 | 15327215 | |
| esg_C4 | 15320871 | 15333970 | |
| esg_D2 | 15322159 | 15327215 | |
| Unspecified_VT7841 | 15322323 | 15324408 | |
| esg_D6 | 15323002 | 15324419 | |
| esg_enhancerE | 15323767 | 15326060 | |
| Unspecified_VT7842 | 15323817 | 15325903 | |
| esg_D3 | 15324419 | 15327215 | |
| Unspecified_VT7843 | 15325315 | 15327423 | |
| esg_D4 | 15325794 | 15327215 | |
| esg_enhancerD | 15325916 | 15327967 | |
| Unspecified_VT7844 | 15326847 | 15328948 | |
| esg_C3 | 15327209 | 15333970 | |
| Unspecified_VT7845 | 15328340 | 15330493 | |
| Unspecified_VT7846 | 15329877 | 15331995 | |
| esg_enhancerA | 15330561 | 15333825 | |
| esg_C2 | 15330583 | 15333970 | |
| Unspecified_VT7847 | 15331387 | 15333912 | |
| esg_C1 | 15333084 | 15333970 | |
| esg_TRC1 | 15335531 | 15339117 | |
| Unspecified_VT7848 | 15336095 | 15338186 | |
| Unspecified_VT7849 | 15337851 | 15339976 | |
| esg_MLC2 | 15338779 | 15341871 | |
| Unspecified_VT7850 | 15339601 | 15341708 | |
| Unspecified_VT7851 | 15341294 | 15343595 | |
| esg_C3 | 15341640 | 15345135 | |
| Unspecified_VT7852 | 15343095 | 15345278 | |
| Unspecified_VT7853 | 15344788 | 15346976 | |
| esg_C4 | 15344987 | 15348264 | |
| Unspecified_VT7854 | 15346465 | 15348732 | |
| esg_C5 | 15348077 | 15351562 | |
| Unspecified_VT7855 | 15348341 | 15350575 | |

| | | | |
|---|---|---|---|
| esg_C6 | 15351196 | 15354699 | |
| esg_TRC7 | 15354529 | 15357870 | |
| Unspecified_VT7859 | 15355410 | 15357565 | |
| Unspecified_VT7860 | 15357147 | 15359402 | |
| esg_C8 | 15357674 | 15361093 | |
| Unspecified_VT7861 | 15358892 | 15361027 | |
| wor_rescue_construct | 15418805 | 15431697 | |
| wor_CRM3 | 15425360 | 15426360 | |
| Unspecified_VT7902 | 15435173 | 15437335 | |
| Unspecified_VT7903 | 15436919 | 15439092 | |
| Unspecified_VT7904 | 15438740 | 15440874 | |
| Unspecified_VT7905 | 15440460 | 15442591 | |
| Unspecified_VT7906 | 15442272 | 15444485 | |
| Unspecified_VT7909 | 15455224 | 15457347 | |
| Unspecified_VT7910 | 15457005 | 15459117 | |
| Unspecified_VT7912 | 15460415 | 15462625 | |
| Unspecified_VT7913 | 15462254 | 15464335 | |
| sna_VT7914 | 15463994 | 15466101 | |
| sna_DP | 15464050 | 15467442 | |
| sna_1.7-1 | 15464306 | 15464708 | |
| sna_1.7 | 15464306 | 15466102 | |
| sna_1.7-2A | 15464680 | 15464974 | |
| sna_1.7-2 | 15464681 | 15465217 | |
| sna_1.7-2B | 15464975 | 15465217 | |
| sna_1.7-3 | 15465196 | 15465670 | |
| sna_1.7-4 | 15465653 | 15466102 | |
| Unspecified_BT05 | 15465823 | 15467239 | |
| sna_FC_tested_fragment | 15468504 | 15469766 | |
| Unspecified_VT7918 | 15471027 | 15473179 | |
| Unspecified_VT7919 | 15472723 | 15475030 | |
| sna_dl_mel | 15473775 | 15476675 | |
| Unspecified_VT7920 | 15474478 | 15476667 | |
| Unspecified_HC_41 | 15477447 | 15478435 | **primary** |
| sna_Busser_old | 15477992 | 15478504 | |
| sna_0.25 | 15478142 | 15478530 | |
| sna_A0.9 | 15478142 | 15479138 | |
| sna_A1.3 | 15478142 | 15479565 | |
| sna_A1.6 | 15478142 | 15479841 | |
| sna_A2.2 | 15478142 | 15480406 | |
| sna_A2.8 | 15478142 | 15481082 | |
| sna_A6.0 | 15478142 | 15485097 | |
| sna_0.9 | 15478169 | 15479138 | |

| | | | |
|---|---|---|---|
| sna_2.8kb | 15478169 | 15481082 | |
| sna_W0.05 | 15478308 | 15481082 | |
| sna_W0.1 | 15478326 | 15481082 | |
| sna_W0.18 | 15478447 | 15481082 | |
| sna_W0.25 | 15478529 | 15481082 | |
| sna_RP | 15479229 | 15479841 | |
| sna_HR | 15479840 | 15480305 | |
| Unspecified_VT7922 | 15480180 | 15482932 | |
| sna_ChIP-19 | 15481617 | 15483118 | |
| Unspecified_sna_enhancer[FL] | 15482063 | 15482575 | |
| sna_shadow_FLL | 15485359 | 15486921 | **shadow** |
| sna_ShadowEnhancer | 15485548 | 15486684 | |
| sna_Odz01 | 15485570 | 15487571 | |
| Unspecified_VT7926 | 15488660 | 15490771 | |

**SUPPLEMENTARY TABLE 2. Genomic position of the barcodes used in this study.**

| **Low-Res *doc*-TAD (chromosome 3L)** | | | | | | |
|---|---|---|---|---|---|---|
| Number | Readout | Coordinate (dm3) | | Center | Size (bp) | Number of oligos |
| | | Start | End | | | |
| 1 | RT1 | 8882600 | 8886000 | 8884300 | 3400 | 45 |
| 2 | RT2 | 8901400 | 8904000 | 8902700 | 2600 | 45 |
| 3 | RT3 | 8918000 | 8922000 | 8920000 | 4000 | 45 |
| 4 | RT22 | 8945000 | 8949000 | 8947000 | 4000 | 45 |
| 5 | RT_fiducial | 8959500 | 8963000 | 8961250 | 3500 | 45 |
| 6 | RT24 | 8972700 | 8976500 | 8974600 | 3800 | 45 |
| 7 | RT7 | 8989000 | 8992600 | 8990800 | 3600 | 45 |
| 8 | RT8 | 9005500 | 9009100 | 9007300 | 3600 | 45 |
| 9 | RT9 | 9020600 | 9024000 | 9022300 | 3400 | 45 |
| 10 | RT10 | 9034500 | 9039000 | 9036750 | 4500 | 45 |
| 11 | RT11 | 8891000 | 8894500 | 8892750 | 3500 | 45 |
| 12 | RT12 | 8909400 | 8913200 | 8911300 | 3800 | 45 |
| 13 | RT13 | 8927000 | 8931000 | 8929000 | 4000 | 45 |
| 14 | RT14 | 8936000 | 8939800 | 8937900 | 3800 | 45 |
| 15 | RT15 | 8952200 | 8955500 | 8953850 | 3300 | 45 |
| 16 | RT16 | 8966500 | 8969500 | 8968000 | 3000 | 45 |
| 17 | RT17 | 8978000 | 8981700 | 8979850 | 3700 | 45 |
| 18 | RT18 | 8995500 | 8999200 | 8997350 | 3700 | 45 |
| 19 | RT19 | 9011000 | 9014500 | 9012750 | 3500 | 45 |
| 20 | RT20 | 9027500 | 9031000 | 9029250 | 3500 | 45 |

| **High-Res *doc*-TAD (chromosome 3L)** | | | | | | |
|---|---|---|---|---|---|---|
| Number | Readout | Coordinate (dm3) | | Center | Size (bp) | Number of oligos |
| | | Start | End | | | |
| 1 | RT25 | 8974562 | 8977955 | 8976258 | 3393 | 50 |
| 2 | RT26 | 8977958 | 8981958 | 8979958 | 4000 | 50 |
| 3 | RT27 | 8982073 | 8985954 | 8984013 | 3881 | 50 |
| 4 | RT28 | 8985957 | 8988814 | 8987385 | 2857 | 50 |
| 5 | RT29 | 8988874 | 8992812 | 8990843 | 3938 | 50 |
| 6 | RT30 | 8992815 | 8995836 | 8994325 | 3021 | 50 |
| 7 | RT31 | 8995839 | 8999945 | 8997892 | 4106 | 50 |
| 8 | RT32 | 9000067 | 9003457 | 9001762 | 3390 | 50 |
| 9 | RT33 | 9003460 | 9006983 | 9005221 | 3523 | 50 |
| 10 | RT34 | 9007073 | 9010836 | 9008954 | 3763 | 50 |
| 11 | RT35 | 9010839 | 9014616 | 9012727 | 3777 | 50 |
| 12 | RT36 | 9014655 | 9018437 | 9016546 | 3782 | 50 |
| 13 | RT37 | 9018440 | 9022234 | 9020337 | 3794 | 50 |
| 14 | RT38 | 9022243 | 9025656 | 9023949 | 3413 | 50 |
| 15 | RT39 | 9025809 | 9030194 | 9028001 | 4385 | 50 |
| 16 | RT40 | 9030253 | 9032999 | 9031626 | 2746 | 50 |
| 17 | RT41 | 9033037 | 9038920 | 9035978 | 5883 | 50 |
| 18 | RT_fiducial | 7882692 | 7909948 | 7896320 | 27256 | 352 |

| High-Res *sna*-TAD (chromosome 2L) | | | | | | |
|---|---|---|---|---|---|---|
| Number | Readout | Coordinate (dm3) | | Center | Size (bp) | Number of oligos |
| | | Start | End | | | |
| 1 | 42 | 15244516 | 15249150 | 15246833 | 4634 | 50 |
| 2 | 43 | 15249210 | 15252570 | 15250890 | 3360 | 50 |
| 3 | 44 | 15252625 | 15256548 | 15254586 | 3923 | 50 |
| 4 | 45 | 15256666 | 15259497 | 15258081 | 2831 | 50 |
| 5 | 46 | 15259613 | 15263050 | 15261331 | 3437 | 50 |
| 6 | 47 | 15263097 | 15266713 | 15264905 | 3616 | 50 |
| 7 | 48 | 15266769 | 15269911 | 15268340 | 3142 | 50 |
| 8 | 49 | 15269955 | 15273483 | 15271719 | 3528 | 50 |
| 9 | 50 | 15273597 | 15277751 | 15275674 | 4154 | 50 |
| 10 | 51 | 15277833 | 15281905 | 15279869 | 4072 | 50 |
| 11 | 52 | 15282022 | 15285682 | 15283852 | 3660 | 50 |
| 12 | 53 | 15285726 | 15290276 | 15288001 | 4550 | 50 |
| 13 | 54 | 15290512 | 15294382 | 15292447 | 3870 | 50 |
| 14 | 55 | 15294503 | 15298220 | 15296361 | 3717 | 50 |
| 15 | 56 | 15298264 | 15301875 | 15300069 | 3611 | 50 |
| 16 | 57 | 15301919 | 15306802 | 15304360 | 4883 | 50 |
| 17 | 58 | 15307021 | 15311482 | 15309251 | 4461 | 50 |
| 18 | 59 | 15311625 | 15316267 | 15313946 | 4642 | 50 |
| 19 | 60 | 15316370 | 15320809 | 15318589 | 4439 | 50 |
| 20 | 61 | 15321011 | 15324567 | 15322789 | 3556 | 50 |
| 21 | 62 | 15324611 | 15328328 | 15326469 | 3717 | 50 |
| 22 | 63 | 15328497 | 15333229 | 15330863 | 4732 | 50 |
| 23 | 64 | 15333288 | 15337082 | 15335185 | 3794 | 50 |
| 24 | 65 | 15337126 | 15341650 | 15339388 | 4524 | 50 |

| | | | | | | |
|---|---|---|---|---|---|---|
| 25 | 66 | 15341694 | 15345889 | 15343791 | 4195 | 50 |
| 26 | 67 | 15345959 | 15350251 | 15348105 | 4292 | 50 |
| 27 | 68 | 15350295 | 15355117 | 15352706 | 4822 | 50 |
| 28 | 69 | 15355161 | 15359653 | 15357407 | 4492 | 50 |
| 29 | 70 | 15359697 | 15363607 | 15361652 | 3910 | 50 |
| 30 | 71 | 15363723 | 15367431 | 15365577 | 3708 | 50 |
| 31 | 72 | 15367507 | 15372341 | 15369924 | 4834 | 50 |
| 32 | 73 | 15372385 | 15376313 | 15374349 | 3928 | 50 |
| 33 | 74 | 15376357 | 15380827 | 15378592 | 4470 | 50 |
| 34 | 75 | 15380881 | 15385217 | 15383049 | 4336 | 50 |
| 35 | 76 | 15385261 | 15388932 | 15387096 | 3671 | 50 |
| 36 | 77 | 15389031 | 15393393 | 15391212 | 4362 | 50 |
| 37 | 78 | 15393437 | 15397578 | 15395507 | 4141 | 50 |
| 38 | 79 | 15397636 | 15401837 | 15399736 | 4201 | 50 |
| 39 | 80 | 15401894 | 15406076 | 15403985 | 4182 | 50 |
| 40 | 81 | 15406187 | 15410495 | 15408341 | 4308 | 50 |
| 41 | 82 | 15410541 | 15414485 | 15412513 | 3944 | 50 |
| 42 | 83 | 15414539 | 15418187 | 15416363 | 3648 | 50 |
| 43 | 84 | 15418246 | 15422459 | 15420352 | 4213 | 50 |
| 44 | 85 | 15422503 | 15426235 | 15424369 | 3732 | 50 |
| 45 | 86 | 15426319 | 15429940 | 15428129 | 3621 | 50 |
| 46 | 87 | 15430031 | 15433872 | 15431951 | 3841 | 50 |
| 47 | 88 | 15433939 | 15437685 | 15435812 | 3746 | 50 |
| 48 | 89 | 15437729 | 15441698 | 15439713 | 3969 | 50 |
| 49 | 90 | 15441742 | 15445265 | 15443503 | 3523 | 50 |
| 50 | 91 | 15445316 | 15456773 | 15451044 | 11457 | 50 |
| 51 | 92 | 15456817 | 15460560 | 15458688 | 3743 | 50 |
| 52 | 93 | 15460604 | 15464621 | 15462612 | 4017 | 50 |

| | | | | | | |
|---|---|---|---|---|---|---|
| 53 | 94 | 15464707 | 15468827 | 15466767 | 4120 | 50 |
| 54 | 95 | 15468871 | 15472655 | 15470763 | 3784 | 50 |
| 55 | 96 | 15472699 | 15476808 | 15474753 | 4109 | 50 |
| 56 | 97 | 15476871 | 15481293 | 15479082 | 4422 | 50 |
| 57 | 98 | 15481373 | 15485768 | 15483570 | 4395 | 50 |
| 58 | 99 | 15485841 | 15489710 | 15487775 | 3869 | 50 |
| 59 | 100 | 15489809 | 15493729 | 15491769 | 3920 | 50 |
| 60 | 101 | 15493915 | 15498235 | 15496075 | 4320 | 50 |
| 61 | 102 | 15498279 | 15502064 | 15500171 | 3785 | 50 |
| 62 | 103 | 15502117 | 15505704 | 15503910 | 3587 | 50 |
| 63 | 104 | 15505748 | 15509179 | 15507463 | 3431 | 50 |
| 64 | 105 | 15509223 | 15513552 | 15511387 | 4329 | 50 |
| 65 | 106 | 15513647 | 15517300 | 15515473 | 3653 | 50 |

**SUPPLEMENTARY TABLE 3. Primers for library amplification used in this study.**

| **Low-Res library** | |
|---|---|
| Name | Sequence (5’ >>> 3’) |
| BB297 (Primer forward) | GACTGGTACTCGCGTGACTTG |
| BB299 (Primer reverse) | GTAGGGACACCTCTGGACTGG |
| T7+BB299 (T7 promoter + Primer reverse) | TAATACGACTCACTATAGGGTGTAGGGACACCTCTGGACTGG |

| **High-Res libraries** | |
|---|---|
| Name | Sequence (5’ >>> 3’) |
| BB193 (Primer forward) | TTGATCTCGCTGGATCGTTCTGCAATG |
| BB280 (Primer reverse) | GGGAGTAGGGTCCTTTGTGTG |
| T7+BB280 (T7 promoter + Primer reverse) | TAATACGACTCACTATAGGGTGGGAGTAGGGTCCTTTGTGTG |

**SUPPLEMENTARY TABLE 4. Sequence of imaging (io) and adapter oligos.**

| **Low-Res library** | |
|---|---|
| Name | Sequence (5' >>> 3') |
| io_RT1 | CACACGCTCTTCCGTTCTATGCGACGTCGGTG/iThioMC6-D//3AlexF647N/ |
| io_RT2 | GCGATCATTGGGCACAACGTCCGCTCTTGGTC/iThioMC6-D//3AlexF647N/ |
| io_RT3 | GACCAAGAGCGGACGTTGTGCCCAATGATCGC/iThioMC6-D//3AlexF647N/ |
| io_RT22 | AGAACGATCCAGCGAGATCAAGTGGAGCTGCG/iThioMC6-D//3AlexF647N/ |
| io_fiducial | CATTGCCGTATGGGCTAGGATGACCTGGCTCG/3RhodRd-XN/ |
| io_RT24 | GCATTCACCCTTGCACGATACCGAGCCACACC/iThioMC6-D//3AlexF647N/ |
| io_RT7 | CGCAACGCTTGGGACGGTTCCAATCGGATC/iThioMC6-D//3AlexF647N/ |
| io_RT8 | CGAATGCTCTGGCCTCGAACGAACGATAGC/iThioMC6-D//3AlexF647N/ |
| io_RT9 | ACAAATCCGACCAGATCGGACGATCATGGG/iThioMC6-D//3AlexF647N/ |
| io_RT10 | CAAGTATGCAGCGCGATTGACCGTCTCGTT/iThioMC6-D//3AlexF647N/ |
| io_RT11 | AAGTCGTACGCCGATGCGCAGCAATTCACT/iThioMC6-D//3AlexF647N/ |
| io_RT12 | CGAAACATCGGCCACGGTCCCGTTGAACTT/iThioMC6-D//3AlexF647N/ |
| io_RT13 | ACGAATCCACCGTCCAGCGCGTCAAACAGA/iThioMC6-D//3AlexF647N/ |
| io_RT14 | CGCGAAATCCCCGTAACGAGCGTCCCTTGC/iThioMC6-D//3AlexF647N/ |
| io_RT15 | GCATGAGTTGCCTGGCGTTGCGACGACTAA/iThioMC6-D//3AlexF647N/ |
| io_RT16 | CCGTCGTCTCCGGTCCACCGTTGCGCTTAC/iThioMC6-D//3AlexF647N/ |
| io_RT17 | GGCCAATGGCCCAGGTCCGTCACGCAATTT/iThioMC6-D//3AlexF647N/ |
| io_RT18 | TTGATCGAATCGGAGCGTAGCGGAATCTGC/iThioMC6-D//3AlexF647N/ |
| io_RT19 | CGCGCGGATCCGCTTGTCGGGAACGGATAC/iThioMC6-D//3AlexF647N/ |
| io_RT20 | GCCTCGATTACGACGGATGTAATTCGGCCG/iThioMC6-D//3AlexF647N/ |

| **High-Res libraries** | |
|---|---|
| Name | Sequence (5’ >>> 3’) |
| adapter_RT25 | ACCGAGCAGGTTAGTTGACGatgatccacgCACCGACGTCGCATAGAACGGAAGAGCGTGTG |
| adapter_RT26 | ACGCTCGACAAGAACAGGACtagacgcaccCACCGACGTCGCATAGAACGGAAGAGCGTGTG |
| adapter_RT27 | CGGCCGGTTGGTCTACGGATactcgtttaaCACCGACGTCGCATAGAACGGAAGAGCGTGTG |
| adapter_RT28 | GTCTGGAACGACGGATTCAGgggtactcgcCACCGACGTCGCATAGAACGGAAGAGCGTGTG |
| adapter_RT29 | CGGTGCGGTGGGATGGATAAaacatcggatCACCGACGTCGCATAGAACGGAAGAGCGTGTG |
| adapter_RT30 | CGTTCGTACCGCGTACTTCGagacgacgcaCACCGACGTCGCATAGAACGGAAGAGCGTGTG |
| adapter_RT31 | AGTGCGCACGAGTTGAACTGtttgctcgcaCACCGACGTCGCATAGAACGGAAGAGCGTGTG |
| adapter_RT32 | TCGCCGCGTTTAGACGGGCTccaatgtaccCACCGACGTCGCATAGAACGGAAGAGCGTGTG |
| adapter_RT33 | CGCGGCGTGAATATCGCGGCagtttccataCACCGACGTCGCATAGAACGGAAGAGCGTGTG |
| adapter_RT34 | TACGGGCCCAGACGTTTCATgctacagcgtCACCGACGTCGCATAGAACGGAAGAGCGTGTG |
| adapter_RT35 | GTGTCCGCCAGTACCGTGAGtttatcgtgcCACCGACGTCGCATAGAACGGAAGAGCGTGTG |
| adapter_RT36 | CGATTAGCGCTCGTGCGCGAttaggtccggCACCGACGTCGCATAGAACGGAAGAGCGTGTG |
| adapter_RT37 | CCGTATCCCTGGCGCGGACTcgtgcgggaaCACCGACGTCGCATAGAACGGAAGAGCGTGTG |
| adapter_RT38 | ACGTAAGGCAGCTTGCGTTAaatccggcgtCACCGACGTCGCATAGAACGGAAGAGCGTGTG |
| adapter_RT39 | CCGTCGCCACGCGGACGCAAgggcgtataaCACCGACGTCGCATAGAACGGAAGAGCGTGTG |
| adapter_RT40 | TACGAGCCCTCTTGGACGGGgcgggattcgCACCGACGTCGCATAGAACGGAAGAGCGTGTG |
| adapter_RT41 | ACGCGCCTTTTACTTAATCGagaagatcgcCACCGACGTCGCATAGAACGGAAGAGCGTGTG |
| adapter_RT42 | CTCGGAGCGTTACTGCGGGCctttgttcggCACCGACGTCGCATAGAACGGAAGAGCGTGTG |
| adapter_RT43 | ATCGGGCCCTTTTGTCTGACtaattccggtCACCGACGTCGCATAGAACGGAAGAGCGTGTG |
| adapter_RT44 | CGTCGCGATACTGGTGTAAGcgtgagaatgCACCGACGTCGCATAGAACGGAAGAGCGTGTG |
| adapter_RT45 | TGACTCGGCTCAGTCGCGGCatcggaacggCACCGACGTCGCATAGAACGGAAGAGCGTGTG |
| adapter_RT46 | GCCGGACCGTGATGCCGTGTtctcatggtcCACCGACGTCGCATAGAACGGAAGAGCGTGTG |
| adapter_RT47 | CGGCGTCGGTAGGCCTTCGCttacgaaggtCACCGACGTCGCATAGAACGGAAGAGCGTGTG |
| adapter_RT48 | GACGGCAAGAGAGCGTGCGTgccatggtacCACCGACGTCGCATAGAACGGAAGAGCGTGTG |
| adapter_RT49 | GTCCCGGAAGCACGGGCGACcggattggtcCACCGACGTCGCATAGAACGGAAGAGCGTGTG |
| adapter_RT50 | GGATTCCGTAGGCACGCCGAgcaccggtcgCACCGACGTCGCATAGAACGGAAGAGCGTGTG |

| | |
|---|---|
| adapter_RT51 | CGTCGTGGGACGTGGACCGTacacccgataCACCGACGTCGCATAGAACGGAAGAGCGTGTG |
| adapter_RT52 | CGCAATCGGAGAACTACACCcgacgatgatCACCGACGTCGCATAGAACGGAAGAGCGTGTG |
| adapter_RT53 | ATCGCCTCAATAAAGGCGACctttacgcggCACCGACGTCGCATAGAACGGAAGAGCGTGTG |
| adapter_RT54 | AAGCGGGCGCACCCGCGCGCtgacatcgatCACCGACGTCGCATAGAACGGAAGAGCGTGTG |
| adapter_RT55 | CGTAGCTCACCCGGTTAGCGgatcgaccggCACCGACGTCGCATAGAACGGAAGAGCGTGTG |
| adapter_RT56 | CGTATGAGGACGCGTCAATGggcgggacacCACCGACGTCGCATAGAACGGAAGAGCGTGTG |
| adapter_RT57 | ACTCTATCGGCATGGAGGTAgggcgacgggCACCGACGTCGCATAGAACGGAAGAGCGTGTG |
| adapter_RT58 | GTCCGTAGGCAAAGGGTCCGagtctgggtcCACCGACGTCGCATAGAACGGAAGAGCGTGTG |
| adapter_RT59 | GACAATGGTGCGAAAGACCCgccatgcgtcCACCGACGTCGCATAGAACGGAAGAGCGTGTG |
| adapter_RT60 | GATGATCCGCTGAAGTCAAAgtgcgcgttaCACCGACGTCGCATAGAACGGAAGAGCGTGTG |
| adapter_RT61 | CGCGAGGAATTGGCGCATCGttgagcgttgCACCGACGTCGCATAGAACGGAAGAGCGTGTG |
| adapter_RT62 | AAGATGCCGTGAGCCTTTCAgcgcggtcccCACCGACGTCGCATAGAACGGAAGAGCGTGTG |
| adapter_RT63 | CTTCCAATCCCTAAGGCCACaccgaatcggCACCGACGTCGCATAGAACGGAAGAGCGTGTG |
| adapter_RT64 | TTCGCCATCGGTCCAAGCTCgcctcccgatCACCGACGTCGCATAGAACGGAAGAGCGTGTG |
| adapter_RT65 | AACTTAGCGATCACGCGCAAcggatcgcccCACCGACGTCGCATAGAACGGAAGAGCGTGTG |
| adapter_RT66 | CTATCGCTGATTCGTCCGGCgaaaccggcaCACCGACGTCGCATAGAACGGAAGAGCGTGTG |
| adapter_RT67 | AGCCGCGAATCGGCCGTCTAatgcgatgagCACCGACGTCGCATAGAACGGAAGAGCGTGTG |
| adapter_RT68 | AAATCCCGGAAATAACGGCCcgccttaccgCACCGACGTCGCATAGAACGGAAGAGCGTGTG |
| adapter_RT69 | AATGGGCATTCGATGCGCAGgcaacgggcaCACCGACGTCGCATAGAACGGAAGAGCGTGTG |
| adapter_RT70 | CGTCAACCACGGCGCTGCAAtaggcagcgaCACCGACGTCGCATAGAACGGAAGAGCGTGTG |
| adapter_RT71 | CCAATGAAGCAACGCGCTTTcagcaggcgtCACCGACGTCGCATAGAACGGAAGAGCGTGTG |
| adapter_RT72 | CGCAAGATGGTGTACGTCCAcgtacccgctCACCGACGTCGCATAGAACGGAAGAGCGTGTG |
| adapter_RT73 | CTTTGTCGCGCCTTACACCAtgcgggtgcgCACCGACGTCGCATAGAACGGAAGAGCGTGTG |
| adapter_RT74 | ATTCACGCGACCGCTTGACAgcgtcagccgCACCGACGTCGCATAGAACGGAAGAGCGTGTG |
| adapter_RT75 | GAGAAGCGCTCGGGTATGACgcgtcgatcgCACCGACGTCGCATAGAACGGAAGAGCGTGTG |
| adapter_RT76 | GTCAACAACGGCGAATCGCTggatgcgccgCACCGACGTCGCATAGAACGGAAGAGCGTGTG |
| adapter_RT77 | TGAAAGCCGGACAGTTCGCAtcggtcaaccCACCGACGTCGCATAGAACGGAAGAGCGTGTG |
| adapter_RT78 | CACTCATCCGGACGACCTCAtcgcgttgacCACCGACGTCGCATAGAACGGAAGAGCGTGTG |
| adapter_RT79 | CACGATAGGCCCATCGCGAAgttgttcacgCACCGACGTCGCATAGAACGGAAGAGCGTGTG |

| | |
|---|---|
| adapter_RT80 | GTCAAGGACGGCAGTGCAAAacaaccggcaCACCGACGTCGCATAGAACGGAAGAGCGTGTG |
| adapter_RT81 | TCGGAGTCAGCCTCGCGACTtcgcttgtgtCACCGACGTCGCATAGAACGGAAGAGCGTGTG |
| adapter_RT82 | TTGATACACATCACTCGCAAcaaggccgtgCACCGACGTCGCATAGAACGGAAGAGCGTGTG |
| adapter_RT83 | ACGCGTCGCCACACGCAATGgttcctctcaCACCGACGTCGCATAGAACGGAAGAGCGTGTG |
| adapter_RT84 | CTTCACCGCGTTTGTTCAACcggtgccaagCACCGACGTCGCATAGAACGGAAGAGCGTGTG |
| adapter_RT85 | CCAGAGTCGGACATCTCCCAtgccgattgcCACCGACGTCGCATAGAACGGAAGAGCGTGTG |
| adapter_RT86 | GACACGTCCCTCAATCGAACtggccgacgcCACCGACGTCGCATAGAACGGAAGAGCGTGTG |
| adapter_RT87 | TTGGACCGGCATGTCAAGAGaccgccgaacCACCGACGTCGCATAGAACGGAAGAGCGTGTG |
| adapter_RT88 | TGGACTTCCGGTCGATATCCgggcgggagtCACCGACGTCGCATAGAACGGAAGAGCGTGTG |
| adapter_RT89 | TGACGTCGGTTTGGACGAGAttcccgccaaCACCGACGTCGCATAGAACGGAAGAGCGTGTG |
| adapter_RT90 | CCCTTGTGAGCGCCCGACATgcgttgatgtCACCGACGTCGCATAGAACGGAAGAGCGTGTG |
| adapter_RT91 | CTCAAGCGGCAATCGTGTTGgttcgggcttCACCGACGTCGCATAGAACGGAAGAGCGTGTG |
| adapter_RT92 | TGTCAGGTCCGCATGGGTCGcgcttatcgaCACCGACGTCGCATAGAACGGAAGAGCGTGTG |
| adapter_RT93 | GTGGATCCCAGAGTGCCAAAaacgtcatcgCACCGACGTCGCATAGAACGGAAGAGCGTGTG |
| adapter_RT94 | TGCTTACGCGACGCACTTGAgcgcttgacaCACCGACGTCGCATAGAACGGAAGAGCGTGTG |
| adapter_RT95 | GTCGTTCCGATCATCGCACAggccagagtgCACCGACGTCGCATAGAACGGAAGAGCGTGTG |
| adapter_RT96 | ACATTGCGTGGACGTACACGcgtccctccgCACCGACGTCGCATAGAACGGAAGAGCGTGTG |
| adapter_RT97 | ACGTCCCGTGTGCTGTCGCGgtccgatgaaCACCGACGTCGCATAGAACGGAAGAGCGTGTG |
| adapter_RT98 | CCAAAGTTGGTCTCGCGCAAgacacgacgtCACCGACGTCGCATAGAACGGAAGAGCGTGTG |
| adapter_RT99 | GACTGCGACGAACTCGATGCatggccacccCACCGACGTCGCATAGAACGGAAGAGCGTGTG |
| adapter_RT100 | CTTCACGACGGTGTCCGCTTcagggcgatcCACCGACGTCGCATAGAACGGAAGAGCGTGTG |
| adapter_RT101 | GCCCACATGCGTAGGCGAGGgaaacatcgtCACCGACGTCGCATAGAACGGAAGAGCGTGTG |
| adapter_RT102 | GCGATTGCCGCTTCAATCAActggcgttgcCACCGACGTCGCATAGAACGGAAGAGCGTGTG |
| adapter_RT103 | AACAAACGGCACACTCGGATggcacgcttcCACCGACGTCGCATAGAACGGAAGAGCGTGTG |
| adapter_RT104 | CCTGCGCACTTGCCCGTGTTtgactcccgaCACCGACGTCGCATAGAACGGAAGAGCGTGTG |
| adapter_RT105 | CCAAGCGCAGACGAACGGCAttcccacacgCACCGACGTCGCATAGAACGGAAGAGCGTGTG |
| adapter_RT106 | CACAAGGTTCGCCACATCTTgccatgaaagCACCGACGTCGCATAGAACGGAAGAGCGTGTG |
| adapter_ fiducial | AATCTTAGCGTGTCCATTGCccgcaggaccAGATCCGGCGGATACACAA |

| | |
|---|---|
| io_RT_all | CACACGCTCTTCCGTTCTATGCGACGTCGGTG/iThioMC6-D//3AlexF647N/ |
| io_fiducial | TTGTGTATCCCGCCGGATCTTTGAATCAAC/3RhodRd-XN/ |

#### SUPPLEMENTARY TABLE 5. Sequence of primary probes

See separate Excel file <Supplementary_Table5.xlsx>.

#### SUPPLEMENTARY TABLE 6. RNA probes information

| Probe | Vector backbone | Restriction enzyme | RNA polymerase |
|---|---|---|---|
| *Sna* | pBluescript II SK (+) | NotI | T7 |
| *Doc1* | pCR II | EcoRV | SP6 |
| *Doc2* | pCR II | SpeI | T7 |
| *Doc3* | pCR II | SpeI | T7 |

| **Primer information** | |
|---|---|
| Primer name | Primer Sequence |
| Doc1_intronic_Forward | CCTATCTCATACGAGTACGAG |
| Doc1_intronic_Reverse | AGGACTTGATTAAGTGGCCTGCC |
| Doc2_Forward | AGCAACTATTGCGTGCTCCT |
| Doc2_Reverse | AAATGGCCGATATGCTGAAG |
| Doc3_Forwad | GTTTACCCGGTAAGTGGAGC |
| Doc3_Reverse | GGCGTTGGTTAATGGATGTT |
| Sna_Forward | ATTTAATTCTTCTCTTTAAGC |
| Sna_Reverse | GGGTAAATCGGGAGATCGGCG |
